# A cryptic tubulin-binding domain links MEKK1 to microtubule remodelling

**DOI:** 10.1101/2020.04.07.030676

**Authors:** Pavel Filipčík, Sharissa L. Latham, Antonia L. Cadell, Catherine L. Day, David R. Croucher, Peter D. Mace

## Abstract

The MEKK1 protein is a pivotal kinase activator of responses to cellular stress. Activation of MEKK1 can trigger various responses, including mitogen activated protein (MAP) kinases, NF-κB signalling, or cell migration. Notably, MEKK1 activity is triggered by microtubule-targeting chemotherapies, amongst other stressors. Here we show that MEKK1 contains a previously unidentified tumour overexpressed gene (TOG) domain. The MEKK1 TOG domain binds to tubulin heterodimers—a canonical function of TOG domains—but is unusual in that it appears alone rather than as part of a multi-TOG array, and has structural features distinct from previously characterised TOG domains. MEKK1 TOG demonstrates a clear preference for binding curved tubulin heterodimers, which exist in soluble tubulin and at sites of microtubule polymerisation and depolymerisation. Mutations disrupting tubulin-binding lead to destabilisation of the MEKK1 protein in cells, and ultimately a decrease in microtubule density at the leading edge of polarised cells. We also show that MEKK1 mutations at the tubulin-binding interface of the TOG domain recur in patient derived tumour sequences, suggesting selective enrichment of tumour cells with disrupted MEKK1–microtubule association. Together, these findings provide a direct link between the MEKK1 protein and tubulin, which is likely to be relevant to cancer cell migration and response to microtubule-modulating therapies.

**SIGNIFICANCE STATEMENT:** The protein kinase MEKK1 activates stress response pathways in response to various cellular stressors, including chemotherapies that disrupt dynamics of the tubulin cytoskeleton. Filipčík et al., show that MEKK1 contains a previously uncharacterised domain that can preferentially bind to the curved tubulin heterodimer—which is found at sites of microtubule assembly and disassembly. Mutations that interfere with MEKK1-tubulin binding disrupt microtubule networks in migrating cells and are enriched in patient-derived tumour sequences. These results suggest that MEKK1-tubulin binding may be relevant to cancer progression, and the efficacy of microtubule-disrupting chemotherapies that require the activity of MEKK1.

## Introduction

Mitogen-activated protein kinase (MAPK) signalling is used throughout eukaryotic organisms as a means of regulating proliferation, differentiation and stress responses. In general, MAPK signalling occurs via hierarchical cascades, where a MAPK is activated following phosphorylation by a MAP kinase kinase (MAP2K), which itself is activated by a MAP kinase kinase kinase (MAP3K). The molecular basis of regulation of MAPKs and MAP2Ks is relatively well-established (Peti and Page, 2013). However, the diversity inherent to the 24 MAP3Ks present in humans means that outside a few well-studied cases, most notably RAF kinases (Kondo et al., 2019; Matallanas et al., 2011; Park et al., 2019), the molecular mechanisms underlying MAP3K regulation remain relatively poorly understood.

MEKK1 (also known as MAP3K1) is unique in that it is the only MAP3K to also possess ubiquitin E3 ligase activity. The ability to phosphorylate or ubiquitinate substrates under different circumstances allows MEKK1 to function as a central signalling hub—it can promote signalling via the ERK, JNK, p38, NF-κB, or other pathways depending on cellular context (Lee et al., 1997, 1998; Xia et al., 1998; Yujiri et al., 1998). Direct targets for MEKK1 kinase activity include: MAP2Ks, leading to activation of JNK, ERK, and p38 pathways; Inhibitor of κ kinases (IκKs), leading to NF-κB transcription factor family activation (Deak and Templeton, 1997; Lee et al., 1998; Xu et al., 1996); or migration-linked proteins such as calponin-3 (Hirata et al., 2016). MEKK1 can ubiquitinate itself in a phosphorylation-dependent manner (Witowsky and Johnson, 2003), leading to attenuation of its own kinase activity, and the diminished activity of downstream ERK and JNK pathways. Alternatively, ubiquitination of MEKK1 can increase its kinase activity upon CD40 stimulation (Matsuzawa et al., 2008). MEKK1 can also ubiquitinate TAB1 with K63-linked polyubiquitin chains, leading to p38, JNK, and TAK1 activation (Charlaftis et al., 2014). MEKK1 is also involved in the control of ERK and c-Jun stability via K48-linked polyubiquitination and proteasomal degradation (Lu et al., 2002; Rieger et al., 2012; Xia et al., 2007).

Activation of MEKK1 can result in many cellular outcomes, ranging from proliferation to apoptosis, depending on cellular context. It is therefore necessary to control its activation both temporally and spatially in order to ensure robust, specific responses to given stimuli. MEKK1 is associated with many subcellular structures, including tight junctions (Steed et al., 2014), actin stress fibers, focal adhesions (Christerson et al., 1999), the cytoplasmic membrane (Gallagher et al., 2007; Matsuzawa et al., 2008), and the perinuclear region (Lu et al., 2002).

In terms of domain architecture, MEKK1 contains N-terminal SWIM-type and RING-type zinc finger domains, and a C-terminal serine-threonine kinase domain (Figure 1A). The SWIM domain is thought to mediate protein-protein interactions and bind c-Jun, while the RING domain exerts ubiquitin E3 ligase activity (Aravind et al., 2003; Charlaftis et al., 2014; Rieger et al., 2012). MEKK1 also contains a caspase-3 cleavage site at Asp-878 that, when cleaved, separates the kinase domain from the SWIM and RING domains (Jarpe et al., 1998; Schlesinger et al., 2002; Tricker et al., 2011). Aside from recruitment to α-actinin and stress fibers via a region incorporating the SWIM-RING domains (Christerson et al., 1999), little is known about the factors that regulate MEKK1 localisation, and thus signalling outcome.

**Figure 1:**
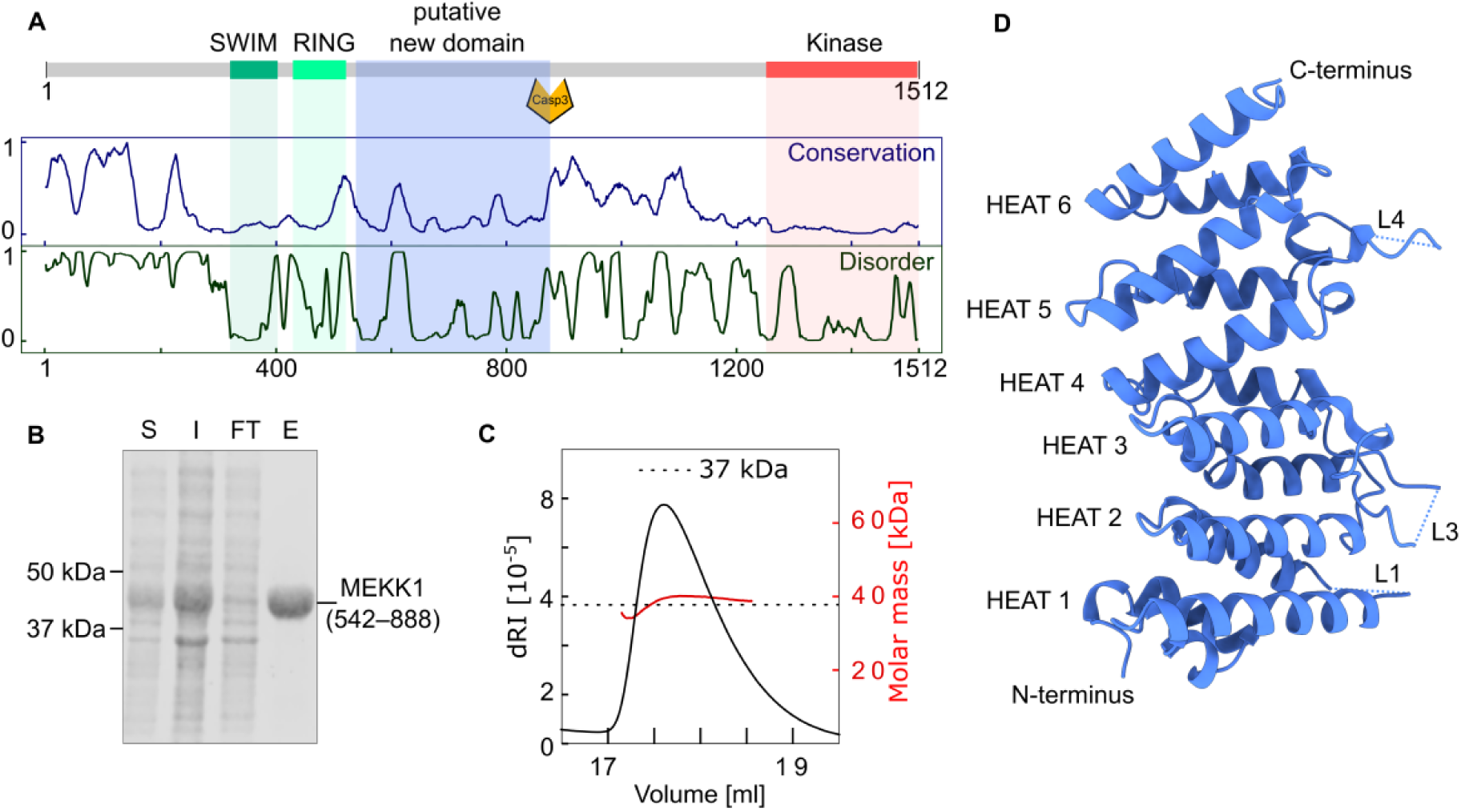
Identification and structure of an uncharacterised domain of MEKK1. A.) Analysis of MEKK1 sequence conservation across species and predicted disorder. Upper panel shows a schematic representation of known features of MEKK1, relative to sequence conservation (middle panel) and predicted disorder (bottom panel). B.) MEKK1 (542– 888) can be expressed and purified from E. coli. S, I, FT, E indicate soluble (S), insoluble (I), flow-through (FT), and eluate (E) fractions from lysis and Ni^2+^ affinity purification, as visualised using Coomassie staining. C.) MALLS analysis of MEKK1(542–888) indicates a monomeric species in solution. D.) The crystal structure of MEKK1(548–867), with termini and structural features labelled. Missing loops are represented by blue dotted lines.

Here, we show that MEKK1 contains a cryptic tumour overexpressed gene (TOG) domain, which allows it to bind directly to tubulin. A crystal structure of the MEKK1 TOG domain reveals notable differences from canonical TOG domains in the α-tubulin binding surface. MEKK1 TOG domain shows clear preference for binding to the curved conformation of tubulin, which is present mainly in soluble tubulin and at dynamic microtubule ends. Disrupting the tubulin-binding surface reduces microtubule density at the leading edge of polarised cells, and has the potential to link microtubule dynamics to MEKK1 activity in chemotherapeutic response.

## Results

### A previously unidentified domain within MEKK1

Since its discovery, MEKK1 has been annotated as containing SWIM, RING and kinase domains (Lu et al., 2002; Rieger et al., 2012; Xu et al., 1996). To gain further insight into the molecular basis for MEKK1 regulation we combined analysis of sequence conservation across species and prediction of protein disorder (Figure 1 A.). From this analysis we observed that a large portion of MEKK1, between the RING domain and the caspase cleavage site, is highly conserved and displays characteristics consistent with the presence of a structured domain. Notably, MEKK1 from diverse metazoans have a conserved region from Asn540 to Gly872 (human numbering), and PONDr (Romero et al., 2001) analysis of the sequence suggested a well-ordered domain (Figure 1 A., Supplementary Figure 1). Because this region had not previously been functionally annotated, we cloned residues 542–888 of human MEKK1 to investigate its function.

Following overexpression of His-tagged protein in *E. coli* (Figure 1 B.), we observed that MEKK1(542–888) eluted as a monodisperse peak upon size-exclusion chromatography. Analysis using size-exclusion chromatography coupled to multiple-angle light scattering (SEC-MALLS) confirmed that the domain was monomeric and that it did not form aggregates in solution (Figure 1 C.). We grew crystals of MEKK1(542–888), solved its structure using experimental phasing (Figure 1 D.; Supplementary Figure 2), and refined the final structure against native diffraction data to a resolution of 1.9 Å (Supplementary Table 1).

### MEKK1 contains a cryptic TOG domain

The structure of MEKK1(542–888) consists of six helix-loop-helix repeats side-by-side in an extended array (Figure 1 D.). Such helix-loop-helix repeats are designated as HEAT (Huntingtin, Elongation factor 3, protein phosphatase 2A, and yeast kinase TOR1) repeats. The loops connecting the helical array showed notable differences in character on either side of the molecule; on one side the loops are compact, well-defined by electron density and conserved in MEKK1 from various organisms, while on the opposing side of the molecule the loops (L1, L3 and L4) are longer, poorly defined by electron density and less well conserved in sequence (Supplementary Figure 3). To assess similarity to known functional domains, we searched for similar structures using the PDBeFold server (Krissinel and Henrick, 2004), which uncovered remarkable similarity to structures of Tumour overexpressed gene (TOG) domains from human, mouse, and drosophila (Supplementary Table 2). In general, TOG domains adopt a similar fold with a core of five HEAT repeats, with some variation at the N- and C-termini. Based on this observed structural similarity, we subsequently refer to the region covering residues 548–867 of MEKK1 as the MEKK1 TOG domain.

The primary role of TOG domains is to mediate direct interaction with tubulin and/or microtubules, which raised the possibility that MEKK1 may bind directly to tubulin. A direct MEKK1-tubulin interaction was particularly intriguing in light of previous evidence that MEKK1 responds to microtubule stress in a manner distinct from its response to other stressors (Gibson et al., 1999; Kwan et al., 2001; Tricker et al., 2011; Yujiri et al., 1999). The canonical function of TOG domains is the regulation of microtubule dynamics through conformation-specific binding to tubulin (Al-Bassam and Chang, 2011; Byrnes and Slep, 2017; Fox et al., 2014; Majumdar et al., 2018; Maki et al., 2015; Slep, 2009). However, modulating microtubule assembly typically requires the presence of two or more TOG domains in an array, and MEKK1 only contains one TOG domain. We therefore set out to ascertain whether the MEKK1 TOG domain does in fact bind to tubulin, and secondly whether binding may be sensitive to tubulin conformation—namely whether it preferentially binds the curved tubulin heterodimer or intact (straight) microtubules.

### Model of a MEKK1-tubulin interaction

In order to better visualize the similarities and differences between the MEKK1 TOG domain and TOG domains that recognise various states of tubulin, a set of models was prepared (Ayaz et al., 2012; Byrnes and Slep, 2017; Kellogg et al., 2017; Nogales et al., 1998). These models allowed direct comparison of the features that drive the conformational preference of different TOG domains relative to the MEKK1 TOG domain (Figure 2). The structures of *S. cerevisiae* Stu2p TOG1 and *D. melanogaster* Msps TOG5 were used for this comparison because they preferentially bind curved and straight tubulin, respectively. The MEKK1 TOG-Stu2p TOG1 alignment has a C_α_ rmsd of 3.84 Å over 185 aligned residues with 13% sequence identity (Figure 2 A. and Supplementary Table 3) and the MEKK1 TOG-Msps TOG5 a C_α_ rmsd of 3.16 Å over 193 aligned residues with 11.4% sequence identity (Figure 2 B. and Supplementary Table 3). Overall, the TOG domain in MEKK1 has strong structural conservation in the core HEAT repeat array with TOG domains that bind either straight or curved tubulin. The main differences between the domains lies in their N- and C-termini, and in the relative orientation of successive HEAT repeats (Figure 2 A. and B.).

**Figure 2:**
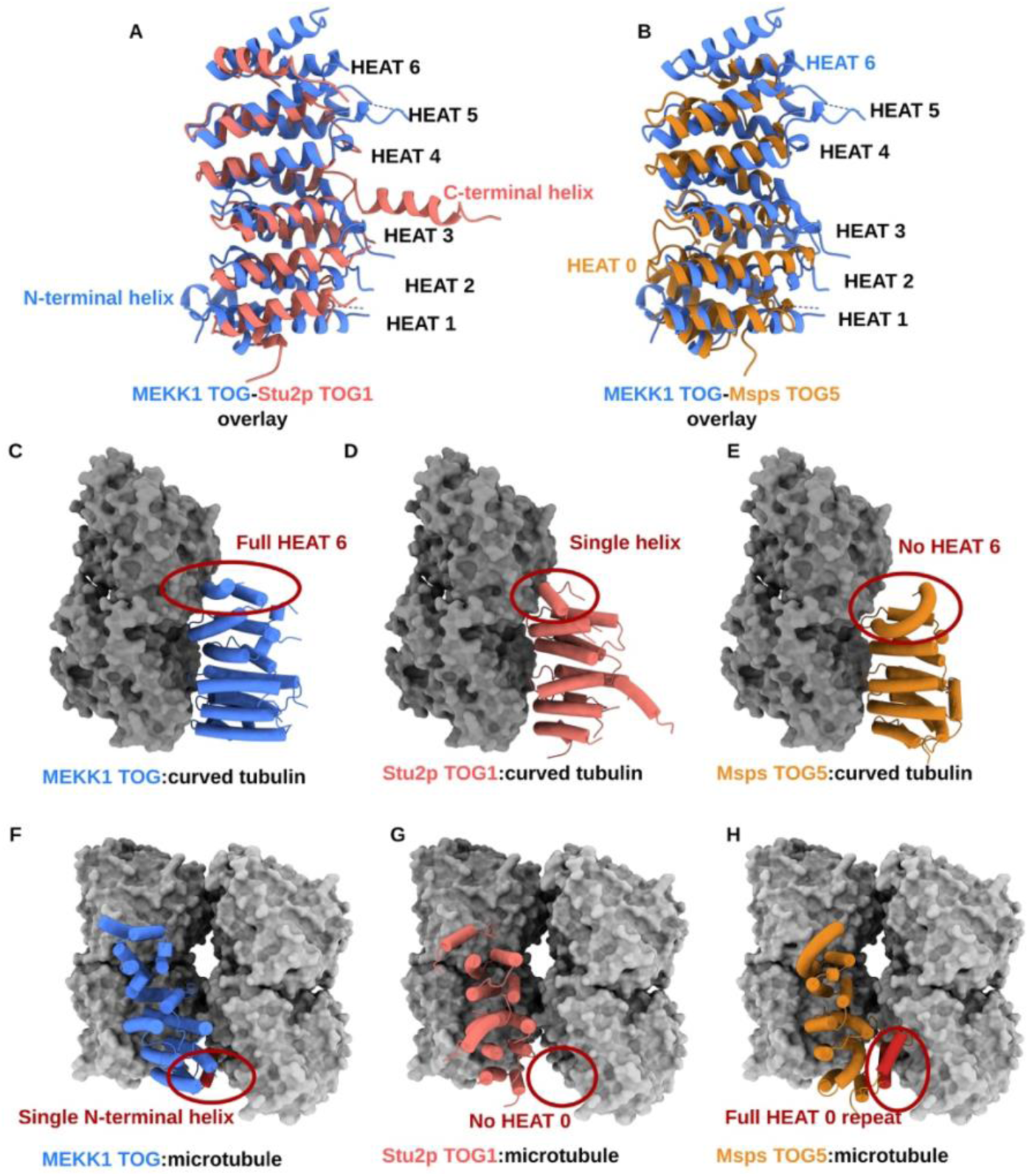
Comparison of tubulin-binding modes of different TOG domains: A.) and B.) Superpositions of the MEKK1 TOG (blue) and Stu2p TOG1 (salmon) or Msps TOG5 (orange), respectively. Shared features are labelled in black and features unique to either structure are labelled in their respective colours. C.) A model of MEKK1 TOG in complex with curved tubulin (PDB ID: 6WHB and 4FFB). D.) Stu2p TOG1 in complex with curved tubulin (PDB ID: 4FFB) E.) A model of Msps TOG5 in complex with curved tubulin (PDB ID: 5VJC and 4FFB) F.) A model of MEKK1 TOG in complex with the microtubule lattice (PDB ID: 6WHB and 5SYF). G.) A model of Stu2p TOG1 in complex with microtubule lattice (PDB ID: 4FFB and 5SYF) H.) A model of Msps TOG5 in complex with microtubule lattice (PDB ID: 5VJC and 5SYF).

Next, we considered the potential of the MEKK1 TOG domain to bind the curved (Figure 2 C.–E.), or straight (Figure 2 F.–H.), forms of tubulin. Overall, the MEKK1 TOG shows electrostatic properties complementary to tubulin across the putative interface (Supplementary Figure 4). When compared to Stu2p, which binds curved tubulin, MEKK1 TOG contains a full C-terminal HEAT repeat where Stu2p has only a single helix (Figure 2 C/D.; red circle). MEKK1 TOG’s additional helix within HEAT 6 is arginine-rich, a feature shared with only one other structurally characterized TOG domain, Cep104 TOG (Supplementary Figure 5). The straight tubulin-binding Msps TOG5 lacks HEAT 6 entirely (Figure 2 E.; red circle). Compared to a straight tubulin binding TOG in Msps, MEKK1 TOG has a single helix at its N-terminus, whereas Msps has a pair of helices known as HEAT 0 (Figure 2 F./H.). HEAT 0 is thought to facilitate microtubule binding by contacting an adjacent protofilament in the microtubule lattice (Al-Bassam et al., 2007; Byrnes and Slep, 2017); Figure 2 H.).

Overall, the MEKK1 TOG domain bears similarities to both curved tubulin-binding TOGs and lattice-binding TOGs (Figure 2 A. and B.), but it is not readily apparent which conformation of tubulin is it likely to bind. Because tubulin conformation-specificity would be crucial to the mechanistic role of MEKK1 TOG, we sought to experimentally characterise tubulin binding by the domain.

### MEKK1 binds tubulin in solution

To investigate whether the TOG domain of MEKK1 is capable of tubulin-binding, we first tested its ability to bind tubulin *in vitro* using size-exclusion chromatography coelution assays with soluble αβ-tubulin. These showed that MEKK1 TOG is able to bind free soluble tubulin (Figure 3 A.–B.), shifting the tubulin dimer to a higher apparent molecular weight. Although this demonstrated an ability to bind the isolated dimer, tubulin can adopt a continuum of conformations in solution (Hsu et al., 2014), so we also evaluated binding to a specific curved conformation of tubulin, stabilised with colchicine (Banerjee et al., 1997; Hsu et al., 2014; Ravelli et al., 2004). Using the same assay we observed that colchicine-stabilised tubulin was still able to form a higher molecular weight complex with the MEKK1 TOG, although slightly less effectively (Figure 3 C. and Supplementary Figure 6). Together, these experiments showed that MEKK1 TOG domain is a *bona fide* TOG domain, and suggested that MEKK1 TOG can bind the curved conformation of tubulin.

**Figure 3:**
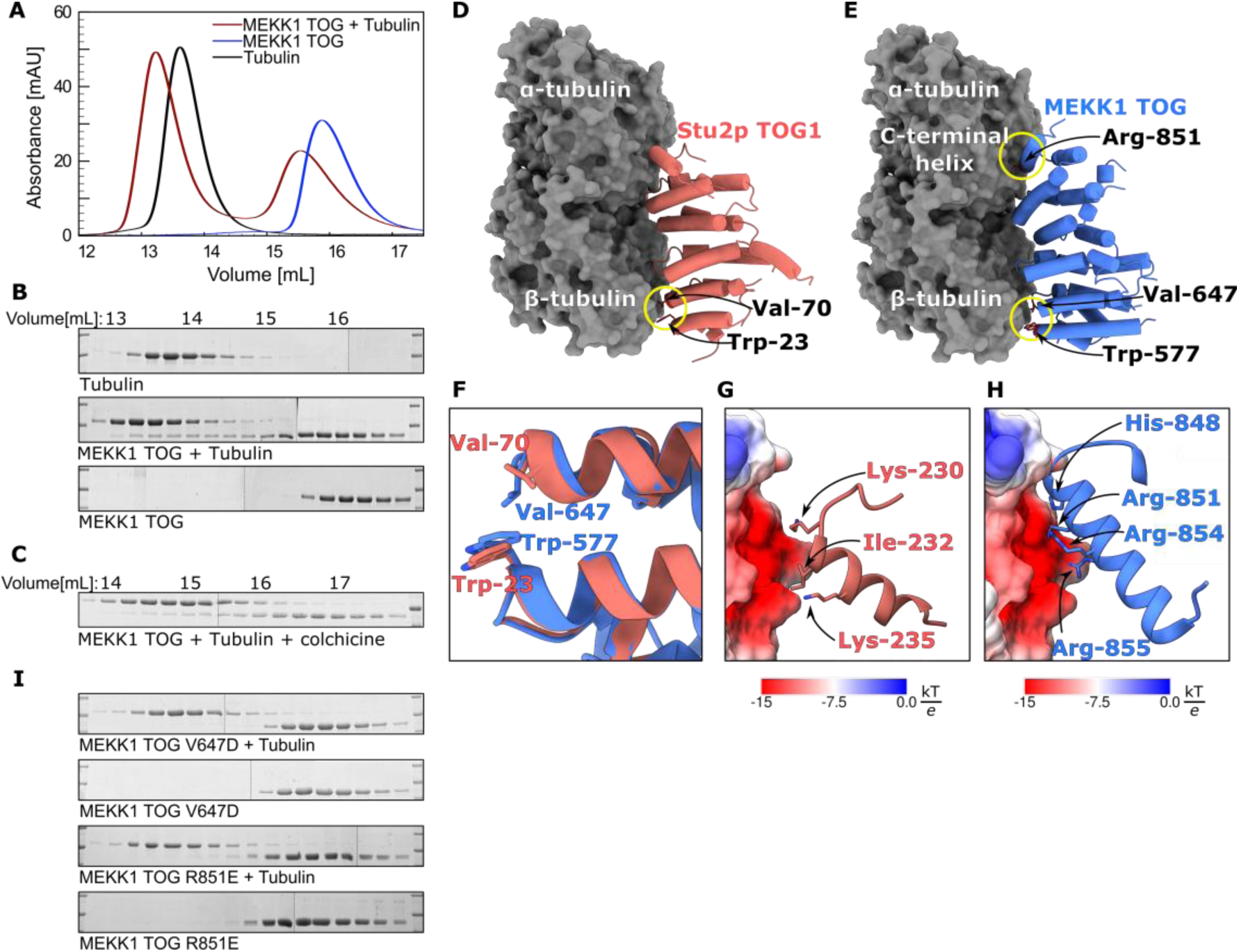
Tubulin binding and mutation analysis of MEKK1 TOG. A.) SEC trace of MEKK1 TOG (blue), tubulin (black), and their combination (red). Earlier elution of TOG and tubulin in combination suggests binding interaction. B.) SDS-PAGE analyses of SEC coelution experiments of MEKK1 TOG with tubulin. C.) SDS-PAGE analysis of SEC coelution assay with MEKK1 TOG and colchicine-treated tubulin. D.) The crystal structure of Stu2p TOG1-tubulin complex, used as the basis for mutant design. E.) A model of MEKK1 TOG interacting with tubulin, highlighting the conserved residues Trp-577 and Val-647, as well as the C-terminal Arg-851, tested in this work. F.) A detail of the β-tubulin interface, conserved between Stu2p TOG and MEKK1. G.) and H.) Details of the C-terminal portions of Stu2p TOG and MEKK1 TOG, respectively, with tubulin surface coloured by electrostatic potential.

We next probed the contribution of different regions of the MEKK1 TOG domain to binding. Although divergent in sequence overall, the β-tubulin contacting residues in Stu2p TOG1 are conserved in MEKK1 TOG (Figure 3 D.–F.). The arginine-rich C-terminal helix of MEKK1 TOG is positioned in the vicinity of the acidic α-tubulin residues, suggesting an extended α-tubulin interaction surface in MEKK1 TOG (Figure 3 G. and H.; Supplementary Figure 4). Thus, we designed mutants at each interface: the β-tubulin interacting Trp-577 and Val-647 patch (Figure 3 F.); and the C-terminal arginine-rich helix of MEKK1 TOG, which significantly differs from the C-terminal helices of Stu2p (Figure 3 G. and H.).

Tubulin coelution assays with MEKK1 TOG mutants showed that both α- and β-tubulin interfaces are involved in the interaction between TOG and tubulin (Figure 3 I., Supplementary Figure 6). On the β-tubulin-interacting side, the Val-647-Asp mutant disrupted tubulin coelution completely (Figure 3 I. and Supplementary Figure 6). The importance of the unique arginine-rich C-terminal HEAT repeat of MEKK1 for tubulin binding was confirmed when a charge swap mutation of Arg-851, situated at the head of the arginine-rich C-terminal helix of MEKK1 TOG (R851E; Figure 3 H.). The R851E mutant disrupted interaction with tubulin to the same extent as a Val-647-Asp mutation in the β-tubulin interface (Figure 3 I.).

These results show that MEKK1 TOG can bind the curved conformation of soluble tubulin, and that both the interfaces implicated by modelling (Figure 2 and 3) are important for the MEKK1 TOG:tubulin interaction.

### Analysis of conformation-specific binding by MEKK1 TOG

As TOG domains can bind tubulin in a conformation-specific manner (Byrnes and Slep, 2017), it was necessary to probe whether the unusual features of the MEKK1 TOG may enable it to recognise straight, as well as curved tubulin. We probed the ability of MEKK1 TOG to bind to the straight conformation of tubulin in the microtubule lattice using paclitaxel-stabilized microtubules in cosedimentation assays (Supplementary Figure 7 A. and B.), negative stain electron microscopy (NS-EM; Supplementary Figure 7 C. and D.), and cryo-electron microscopy (Cryo-EM; Supplementary Figure 7 E. and F.). In cosedimentation assays, a greater proportion of MEKK1 TOG was found in the insoluble fraction cosedimented with paclitaxel-stabilized microtubules than in control experiments lacking microtubules (Supplementary Figure 7 A. and B.). The presence of MEKK1 TOG in the insoluble fraction suggested that interaction with the straight conformation of tubulin may be possible, so this was probed further by electron microscopy. Although NS-EM suggested density on the outside of the microtubule lattice with regular spacings (compare Supplementary Figure 7 C. and D.), MEKK1 TOG binding to paclitaxel-stabilized microtubules was not apparent by cryo-EM. No densities consistent with a microtubule-binding protein were apparent in raw micrographs of microtubules in the presence of MEKK1 TOG (Supplementary Figure 7 E.), and 2D class averages of helix-particles from the cryo-EM preparation show no classes with regularly spaced densities decorating the outside of the microtubule (Supplementary Figure 7 F.).

With cosedimentation assays and electron microscopy studies proving equivocal, we turned to isothermal titration calorimetry (ITC) to test for conformation specific binding. ITC made it possible to test binding to the free and polymerized, paclitaxel-stabilized form of tubulin directly, using the same experimental conditions in both cases. A global analysis of the thermograms of four independent experiments with free tubulin (Figure 5 A., shown in blue, maroon, yellow, and orange) clearly show binding a 1:1 stoichiometry with calculated dissociation constant K_D_ = 0.63 μM. The control titrations of TOG into buffer and buffer into tubulin (Figure 5 A., both shown in cyan) both show negligible contributions of the dilution effects. No interaction was detectable upon injection of MEKK1 TOG domain into paclitaxel-stabilized microtubules (Figure 5 B., shown in blue and maroon).

**Figure 4:**
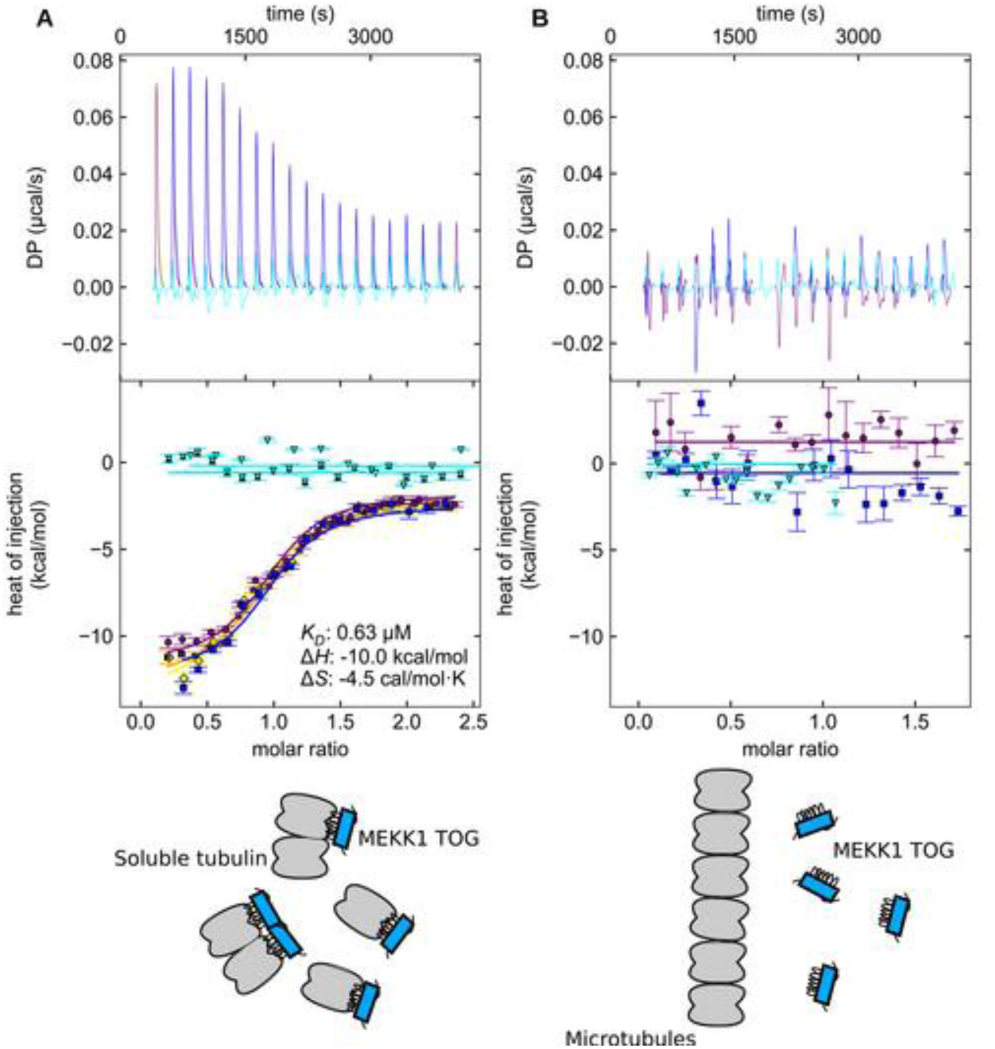
ITC comparison between MEKK1 TOG interacting with the free tubulin and paclitaxel-stabilized microtubules. A.) Thermograms (top) and fitted isotherms (bottom) from a global analysis of four experiments measuring MEKK1 TOG binding to tubulin in non-polymerizing conditions, shown in maroon, orange, yellow, and blue. MEKK1 TOG dilution experiment and tubulin dilution experiment shown in cyan. B.) Thermograms (top) and fitted isotherms (bottom) from two experiments measuring MEKK1 TOG binding to stabilized microtubules, shown in maroon, and blue. Microtubule dilution experiment shown in cyan.

**Figure 5:**
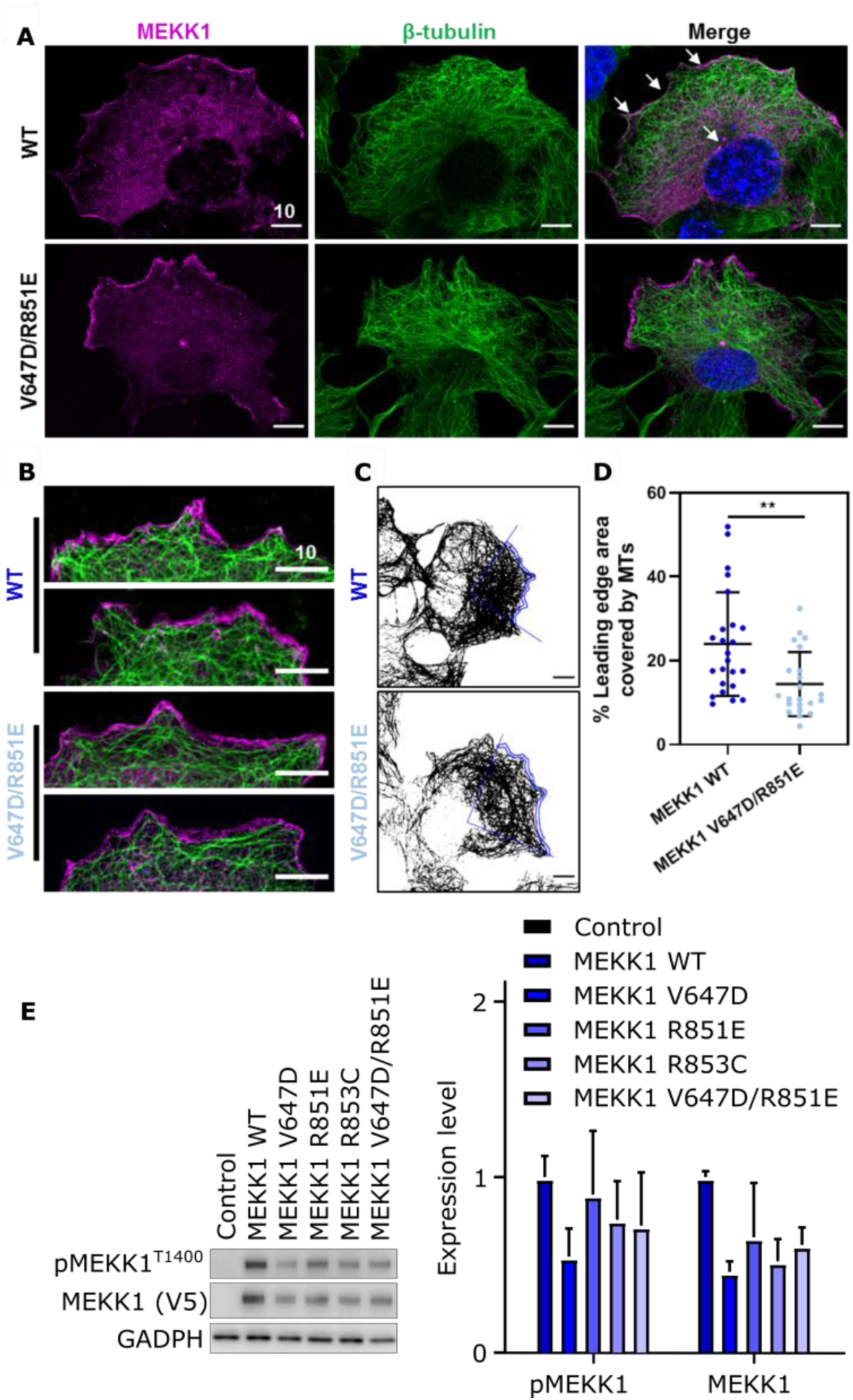
Analysis of mutant MEKK1 in COS-7 cells A.) Distribution of WT (top) and V647D/R851E (bottom) MEKK1 and tubulin in COS-7 cells. MEKK1 signal is shown as magenta, tubulin as green, and DAPI blue. The arrows indicate MEKK1 enrichment at the leading edge and at centrosomes. B.) A detail of the leading edge of polarized cells expressing WT MEKK1 (upper panel) and MEKK1 V647D/R851E (lower panel). C.) An illustration of the quantification of leading edge microtubule density. D.) Fraction of leading edge area covered by microtubules in WT MEKK1-expressing cells (left, dark blue) and MEKK1 V647D/R851E-expressing cells (right, light blue). E.) A representative western blot analysis of the expression and phosphorylation of MEKK1 in HEK-293T cells upon overexpression of WT and mutant MEKK1. Quantification of western blot analyses from three biological replicates, normalised to GADPH levels shown on the right.

Thus, the TOG domain of MEKK1 shows clear preference in binding to the non-polymerized form of tubulin, as opposed to polymerized, paclitaxel-stabilized microtubules. The non-polymerized (curved) form of the tubulin heterodimer is also found at polymerising and depolymerising microtubule ends (Nawrotek et al., 2011; Pecqueur et al., 2012), suggesting that the MEKK1 TOG domain most likely functions to bind tubulin at these sites.

### TOG-tubulin effect on localisation and MAPK activation

Based on a binding preference for curved tubulin—found at sites of microtubule remodelling—we visualised the cellular localisation of wild type MEKK1 and a double mutant incorporating two disruptive mutants at the tubulin binding interface (V647D/R851E). In general, both the wild type and mutant MEKK1 are observed throughout the cell, but are enriched at the cytoplasmic membrane (Figure 5 A.). Remarkably, microtubule density at the cellular periphery visualised by tubulin staining appeared to be reduced in mutant-transfected cells (Figure 5 B.). To quantify this effect, we analysed cells within a scratch assay in order to induce a clear polarized morphology, where the nucleus was positioned towards the rear of the cell and the centrosomes were repositioned towards the cells leading edge. The leading edge was manually demarcated using the MEKK1 signal, and the coverage of microtubules within a 15-pixel region of interest from the leading edge was quantified (Figure 5 C.). In this analysis, coverage of microtubules at the leading edge is significantly reduced in the presence of the MEKK1 TOG V647D/R851E mutant compared to the WT (Figure 5 D.). The effect was independent of the size of the leading-edge region analysed (Supplementary Figure 8). Thus, disrupted tubulin binding by MEKK1 appears to result in less microtubule penetration at the migrating leading edge.

We also probed whether the MEKK1 TOG–tubulin interaction altered the global activation of MAPK pathway activity in cells. To this end, WT full length MEKK1 and a suite of mutants (V647D, R851E, R853C, and V647D/R851E) were expressed in the HEK-293T cell line. In this system, expression of WT full length MEKK1 resulted in the heightened activation of JNK, ERK and p38. While the mutant MEKK1 proteins were generally expressed at lower levels (Figure 5 E.), they still appeared to be capable of activating downstream MAPK pathways. The activation of JNK, ERK, and p38 was reduced across each mutant, but this largely correlated with expression levels of each respective MEKK1 protein (Supplementary Figure 8 B.). One exception was the double mutant V647D/R851E, with mutations in both the α- and β-tubulin binding surfaces of MEKK1 TOG, which showed elevated activation of the p38 pathway.

### MEKK1–Tubulin interface mutants are enriched in tumour samples

Mutations in MEKK1 occur at elevated rates in breast and other cancer types (Cerami et al., 2012; Easton et al., 2007). We therefore analysed whether any of the mutations impact TOG domain function (Cerami et al., 2012; Gao et al., 2013). Nonsense mutations that truncate the protein prior to the C-terminal kinase domain are common in MEKK1— in line with the kinase activity promoting apoptosis. Missense mutations are also prevalent across the gene, with two of the five recurrent sites found within the C-terminal portion of the TOG domain (Figure 6 A.). Modelling the position of these sites (Arg853 and Asp806) on a putative MEKK1-tubulin complex suggests that each lies at the tubulin-binding interface (Figure 6 B.). The non-conservative amino-acid substitutions suggest a high likelihood that they would disrupt tubulin-binding function. In line with this, a R853C mutant displayed markedly decreased stability when ectopically expressed in HEK293T cells (Figure 5 E.). Overall, mutations in the TOG domain of MEKK1, particularly at the tubulin-binding interface, are selectively enriched in a subset of tumour samples, suggesting they may confer a survival advantage in the context of oncogenesis.

**Figure 6:**
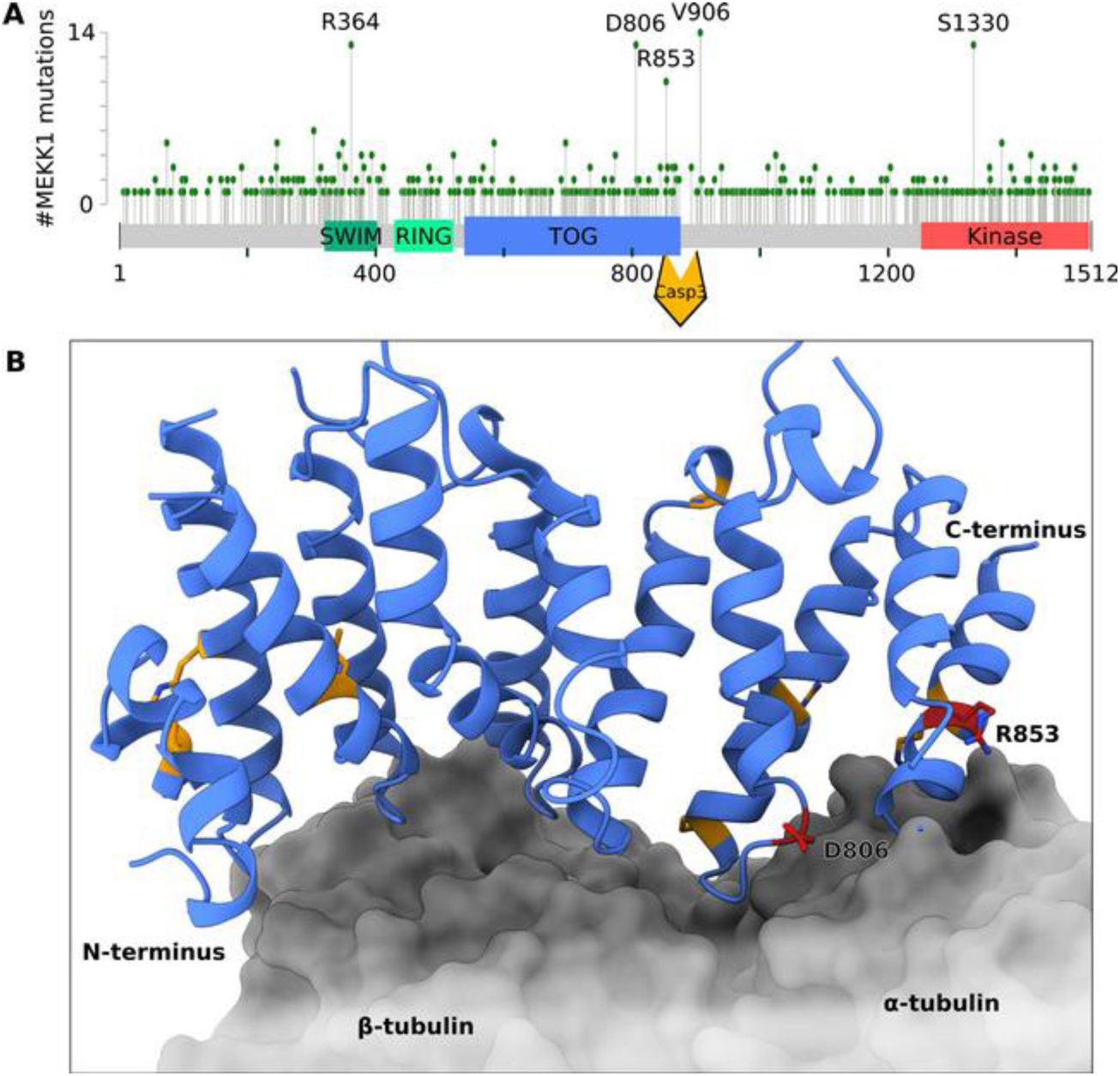
MEKK1 TOG domain harbours mutations in human cancers. A.) Missense mutations in MEKK1 in a curated set of non-redundant human cancer sequence dataset. The TOG domain is one of four mutation hot-spots in the MEKK1 sequence. B.) A model of MEKK1 TOG-tubulin interaction, with MEKK1 TOG domain residues mutated three or more times coloured orange and residues mutated ten or more times coloured red. Asp-806 and Arg-853 both lay on the interface with α-tubulin.

## Discussion

This work presents evidence for the presence of a previously uncharacterised domain within MEKK1, spanning residues 548–867 of the human protein. Based on a crystal structure and structural homology, this domain of MEKK1 was predicted to be a tubulin-binding TOG domain, which was confirmed by *in vitro* binding studies and mutagenesis. ITC experiments show that MEKK1 TOG domain selectively binds the curved conformation of tubulin. The curved conformation of tubulin is associated with non-polymerized tubulin, microtubule end-structures, and microtubule catastrophes. An investigation into the TOG domain’s involvement in MEKK1 in cells suggested that expression of mutant MEKK1 unable to bind tubulin induces less microtubule density at the leading edge of polarised cells. Collectively, these results suggest that the MEKK1 TOG shares some features of canonical TOG domains, while re-purposing the TOG fold to function as a localising agent for the control of signal transduction.

TOG domains are traditionally involved in the regulation of microtubule dynamics. Current models of TOG-catalysed tubulin polymerisation all rely on the presence of multiple TOG domains in a single polypeptide or a complex (Ayaz et al., 2014; Geyer et al., 2018; Nithianantham et al., 2018; Slep and Vale, 2007). MEKK1 only contains one TOG domain, suggesting that the MEKK1 TOG domain mechanism-of-action is different to that of other TOG domains. At the structural level, MEKK1 has common TOG features, but also divergence at the N- and C-termini that may facilitate its role. One close structural homolog of MEKK1 is another recently-identified TOG domain from CEP104 (Supplementary Figure 5; Supplementary Table 3; (Al-Jassar et al., 2017; Rezabkova et al., 2016)). The CEP104 and MEKK1 TOG domains both share a similar pair of additional C-terminal helices relative to canonical TOGs. However, the Cep104 protein is proposed to be multimeric due to oligomerization through an adjacent coiled-coil domain, thus enabling it to function in a similar manner to canonical TOG arrays (Rezabkova et al., 2016).

The role of MEKK1 in mediating the cellular response to stressors that alter cell shape is well-documented. MEKK1 has been found to mediate apoptosis following actin disruption, intermediate filament disruption, and microtubule disruption (Kwan et al., 2001; Tricker et al., 2011), as well as cold shock and hyperosmotic stress, which both induce cytoskeletal remodelling (Breton and Brown, 1998; Nunes et al., 2013; Yujiri et al., 1998). In terms of caspase-dependent activation of MEKK1 (Cardone et al., 1997; Deak et al., 1998; Jarpe et al., 1998; Shiah et al., 2001), the presence of a TOG domain directly N-terminal to the caspase-cleavage site suggests cleavage would release the kinase domain from association with tubulin. In terms of caspase-independent activation of MEKK1 (Kwan et al., 2001; Tricker et al., 2011; Yujiri et al., 1998), the role of tubulin-binding in activation remains to be fully determined. Because it has a much higher affinity for the curved conformation of tubulin, MEKK1 TOG domain could be involved in sensing the status of microtubules directly. Such function would be relevant during microtubule catastrophe—rapid localised depolymerization of microtubules from the straight conformation into the curved conformation—which could bring MEKK1 to the sites of catastrophe and induce clustering of MEKK1.

In unstressed cells expressing mutant MEKK1, we see a defect in the formation of microtubule networks at the leading edge. This finding suggests that MEKK1 may play a role in the crosstalk between dynamic microtubule ends and the actin cytoskeleton, given that it is apparently able to directly bind to both types of filament that are present at the leading edge (Dogterom and Koenderink, 2019). How MEKK1 is regulated and the specific substrates at the leading edge both remain pertinent questions. Logical candidates for interaction partners include: Rho GTPases, previously shown to regulate MEKK1 and actin dynamics (Dogterom and Koenderink, 2019; Gallagher et al., 2004); and Calponin-3, a MEKK1 kinase substrate within the actin cytoskeleton known to impact cellular contractility (Hirata et al., 2016).

Either direct sensing of microtubule catastrophes, or involvement in cellular contractility and migration would be quite relevant to the role of MEKK1 in cancer treatment and metastasis, respectively. Somatic MEKK1 mutations have been identified as significantly enriched in cancer samples in prostate, lung, breast, and ovarian tumour samples (Kan et al., 2010). The finding that two of the most frequent missense mutations in cBioPortal occur at the tubulin-binding interface point to an important role for tubulin association in normal cellular homeostasis, which can be lost in cancer to confer a survival advantage. One possible mechanism would be mutations removing a mechanism of MEKK1 clustering at nascent curved protofilaments, which could serve as transient scaffolds for MEKK1 oligomerization and activation. Alternatively, it could remodel MEKK1’s interactions with other microtubule-associated proteins during migration and metastasis. For instance, ERK1/2 (Harrison and Turley, 2001; Haycock et al., 1992), known microtubule-associated protein kinases and confirmed downstream targets of MEKK1 (Karandikar et al., 2000; Yujiri et al., 1998). However, the exact mechanism by which tubulin-binding mutants impact cancer progression remains to be determined.

MEKK1 is a signalling protein uniquely positioned at the crossroads between ubiquitin and phosphorylation signalling pathways. A direct interaction between MEKK1 and tubulin expands the repertoire of this versatile signalling protein and provides a mechanism for the sensing of microtubule status by MEKK1. Such function could affect substrate selection of MEKK1 kinase or ubiquitin-ligase activities and eventual cellular response. Future studies aimed at uncovering the exact mechanism by which tubulin regulates MEKK1 activity will be relevant both to understanding the role of MEKK1 in normal stress responses, and the efficacy of microtubule disrupting therapies in patients bearing MEKK1 mutations.

## Acknowledgements

This work was supported by Rutherford Discovery Fellowship from the New Zealand government administered by the Royal Society of New Zealand, held by PDM; University of Otago Doctoral Scholarship, held by PF. We also thank the New Zealand synchrotron group for facilitating access to the MX beamlines at the Australian Synchrotron, as well as the beamline staff of the Australian Synchrotron. Structural biology applications used in this project were compiled and configured by SBGrid (Morin et al., 2013).

We are thankful to Rui Zhang and Carolyn Moores for helpful discussions regarding tubulin preparation and electron microscopy.

## Materials and Methods

### Sequence analysis

Q13233 protein sequence from uniprot.org provided a reference protein sequence of MEKK1, and NM_005921.1 mRNA sequence from the NCBI database provided a reference nucleotide sequence of MEKK1. Sequence-based disorder prediction was carried out by PONDr VL-XT (Romero et al., 2001) using the PONDr webserver. ConSurf (Ashkenazy et al., 2010; Celniker et al., 2013; Landau et al., 2005) provided per-residue conservation scores by search with the reference *H. sapiens* MEKK1 sequence in ConSeq mode with default settings.

### Cloning

ProteinCCD was used to aid in design of MEKK1 constructs and primers for ligation-independent cloning (Mooij et al., 2009). MEKK1 542–888 construct (*H. sapiens* MEKK1 TOG) was amplified from the MegaMan human transcriptome library (Agilent). The TOG construct was cloned into pETNKI-his-3C-LIC-kan vector, which was a gift from Titia Sixma (Addgene #108703). Site-directed mutagenesis was carried out by a modified QuickChange protocol using overlapping primers with mutagenic mismatch and 3’ overhangs (Liu and Naismith, 2008).

### Protein purification

All constructs were expressed in E. coli BL21(DE3) in LB media, induced with IPTG overnight at 18 °C, and lysed by sonication. The cell lysate was clarified by centrifugation before protein purification. After purification, all proteins were aliquoted and snap-frozen in liquid nitrogen before storing at –80 °C. Before use, snap-frozen protein aliquots were thawed at room temperature (MEKK1 constructs), or at 37 °C (tubulin and components of the ubiquitin cascade), centrifuged at 20000 g at 4 °C, and kept on ice until use.

The MEKK1 TOG domain was purified for crystallization by Ni^2+^ affinity chromatography, 3c protease cleavage and dialysis against 300 mM NaCl, 10 mM HEPES pH 7.6, 2 mM DTT. For all other purposes, the domain was further purified by size-exclusion chromatography (SEC) in 300 mM NaCl, 10 mM HEPES pH 7.6. Tubulin was purified from fresh bovine brains using two rounds of polymerization-depolymerization in high molarity PIPES buffer (Castoldi and Popov, 2003).

### Crystallization

MEKK1 TOG domain was used for crystallization trials at concentrations of 3.5–30 mg/ml in HBS300 and protein:precipitant drop ratios of 33%–90% in total volumes of 300–1200 nl. Crystallization plates were set up using the mosquito^®^ nanolitre-pipetting system (TTP Labtech) in MRC 2- or 3-well crystallization plates (Swissci) using the high-throughput screens Index™ HT, SaltRx™ HT (Hampton Research) and Morpheus^®^ HT (Molecular Dimensions). Home-made solutions for crystallization screen expansion or cryoprotection were prepared from analytical grade chemicals and filtered through a 22 μm syringe filter before use. The crystal used for native data collection was grown in 1.5 M sodium acetate and 0.1 M BIS-TRIS propane pH 6.9 at a drop ratio of 4:1, and cryoprotected in 1.8 M sodium acetate, 0.1 M BIS-TRIS propane pH 6.9, and 30 % glycerol. The crystal used for anomalous data collection was grown in 1.6 M sodium acetate, 0.1 M BIS-TRIS propane pH 7 at a drop ratio of 9:1, and soaked for 30 seconds in cryoprotectant consisting of 1.8 M sodium acetate, 0.1 M BIS-TRIS propane pH 6.9, 30 % glycerol, and 500 mM sodium iodide before freezing.

### Data collection and structure solution

X-ray diffraction data collection was performed at the Australian Synchrotron using the MX-1 beamline and ADSC Quantum 210r detector (Cowieson et al., 2015). Native dataset was collected at a wavelength of 0.9537 Å and anomalous dataset was collected at a wavelength of 1.4586 Å.

Raw data were processed with XDS (Kabsch, 2010), the space group was determined with Pointless (Evans, 2006, 2011), and the dataset was scaled with Aimless (Evans and Murshudov, 2013). The aimless package also utilizes Ctruncate, Mtzdump, Unique, Freerflag and Cad programs to output the scaled dataset.

The structure was solved using Auto-Rickshaw’s SIRAS pipeline (Panjikar et al., 2005)

### Refinement

The initial model was rebuilt and refined using ARP/wARP (Langer et al., 2008; Morris et al., 2004), followed by iterative manual model modification and refinement using coot (Emsley and Cowtan, 2004), phenix (Adams et al., 2011), and refmac (Murshudov et al., 2011). After initial refinement against the dataset truncated at 2.1 Å resolution with R_work_/R_free_ values of 0.2053/0.2338, the initial dataset was reprocessed with XDS to a resolution of 1.9 Å, including weak high-resolution reflections displaying statistically significant CC1/2. The model refined against the 2.1 Å data was then used for automated building using phenix.autobuild with the 1.9 Å data. Subsequent refinement using coot and phenix.refine yielded a fully refined structure with R_work_/R_free_ of 0.2071/0.2297. A simple composite-omit map was generated using phenix.composite_omit_map. The diffraction data and refined structure coordinates were deposited in the PDB under PDB ID 6WHB.

### Size-exclusion chromatography

In coelution assays, purified MEKK1 constructs were mixed with tubulin in a 2–3:1 molar ratio. DTT was added to a concentration of 1 mM. 100–500 µl of the protein mixture were injected on Superdex 200 Increase column, connected to an ÄKTA pure FPLC system (both GE Healthcare). 250 µl fractions were collected and analysed on SDS-PAGE. In coelution assay with colchicine, tubulin was preincubated with 1 mM colchicine at 0 °C for 20 minutes prior to size-exclusion, and the size-exclusion buffer contained 20 µM colchicine.

### Multi-angle laser light scattering (MALLS)

MALLS data was collected using a Wyatt Dawn 8+ detector (Wyatt Technology) and Waters 410 differential refractometer (Waters) in line with an ÄKTA pure chromatography system fitted with a Superdex 200 Increase column (both GE Healthcare). Data was analysed using ASTRA 5.3.4 software (Wyatt Technology).

### Model preparation

Structural overlays were prepared using secondary structure matching in coot and MatchMaker in UCSF ChimeraX (Krissinel and Henrick, 2004). Source coordinates’ pdb codes are stated in the respective figure legends.

### Isothermal titration calorimetry (ITC)

A low volume Affinity ITC instrument controlled by ITCRun software was used to collect ITC data (both TA instruments). 400 μl of TOG domain at 13.42 mg/ml were desalted into 250 mM NaCl, 250 μM TCEP and 20 mM sodium phosphate at pH 6.5 (ITC buffer) using a 5 mL HiTrap Desalting column (GE Healthcare). Likewise, 80 μl of tubulin at 10.9 mg/ml were desalted into the same buffer. The top fractions from each desalting run were collected, their concentration was determined, and both were degassed at 40 kPa at 25 °C for 10 minutes.

100 mM GTP in water was diluted with ITC buffer to a concentration of 1 mM and 1 μl/100 μl of tubulin was added for a concentration of 10 μM in cell. The sample cell was loaded with 350 μl of tubulin/GTP solution and the ITC syringe was loaded with 320 μl of TOG solution. An ITC run consisted of 1 injection of 0.5 μl, followed by 19 injections of 2.5 μl at 30 °C with a stirring rate of 75 rpm with automatic equilibration set for collection of data with small heats. For each replicate run, tubulin was prepared fresh in the same manner.

For TOG dilution control, tubulin was substituted for the buffer supplemented with GTP as described above. For Tubulin dilution control, the TOG solution was substituted with buffer. The collected data were exported to .xml format using NanoAnalyze (TA Instruments), the baselines were determined using NITPIC, and the thermodynamic parameters of the interaction were determined in a global analysis using SEDPHAT (Brautigam et al., 2016; Zhao et al., 2015). The data were plotted using GUSSI (Brautigam et al., 2016). The blank runs were not subtracted from any of the interaction runs but were plotted alongside the interaction thermograms and isotherms instead.

To probe the interaction between stable microtubules and MEKK1 TOG, stable microtubules were prepared, and their buffer was exchanged by two rounds of centrifugation at 20800 g for 20 minutes and resuspension of the microtubule pellet in ITC buffer. All other actions were done exactly as in the case of probing soluble tubulin-MEKK1 TOG interaction.

### Stable microtubule preparation

Stable microtubules were prepared using paclitaxel stabilization (Moores, 2008). An aliquot of tubulin at 4–10 mg/ml was thawed and mixed with an equal volume of pre-warmed 2 × Pol buffer and incubated the mixture at 37 °C for 10 minutes. Paclitaxel (LC Labs, Woburn, MA, USA) was then added stepwise: 1 μl 100 μM paclitaxel in 5% DMSO was added at the 10- and 15-minute mark, and 1 μl 2 mM paclitaxel in 100% DMSO at the 20- and 25-minute mark. 8 μl of 2 mM paclitaxel in 100% DMSO was added at the 30-minute mark and the microtubules were left to polymerize at 37 °C for another 3 hours. The polymerized microtubules were then transferred to room temperature and left to stabilize for at least 16 hours prior to use. Their concentration was determined by diluting them 10-fold in 5 mM CaCl_2_ in 80 mM PIPES-K pH 6.8, 1 mM MgCl_2_, 1mM EGTA (BRB80), leaving them to depolymerize for at least 30 minutes on ice, measuring A_280_, and calculating concentration with ε = 115000 and M = 110 kDa.

### Cosedimentation assay

The stabilized microtubules were separated from smaller species by centrifugation for 20 minutes at 20800 g at 20 °C and the pellets were resuspended in BRB80 + 20% glycerol. 250 μl BRB80 supplemented with 20% glycerol was added to Eppendorf tubes, a 2:1 molar mixture of TOG and microtubules was layered on top of the glycerol solution and the samples were centrifuged at 20800 g for 40 minutes at room temperature. The top 20 μl were aspirated as the top fraction, the middle 230 μl discarded as the intermediate fraction and 20 μl of 2 × SDS-PAGE buffer was added to the bottom 20 μl and the pellet was resuspended. The contents of the top and bottom fractions were analysed using SDS-PAGE.

### Negative stain electron microscopy

300-mesh in-house carbon-coated copper grids were glow discharged using BioRad E5100 SEM coating system modified in-house for glow discharging grids (BioRad Microscience Ltd, Hertfordshire, UK) in low vacuum (<200 Pa) with a current of 15 mA for 60 seconds.

0.9 mg/ml paclitaxel-stabilized microtubules were mixed with MEKK1 TOG at 5.5 mg/ml, incubated at room temperature for 30 min, and centrifuged at 20800 g for 20 min at room temperature. The supernatant was discarded, the pellet resuspended in 300 mM NaCl in 10 mM HEPES pH 7.6, and immediately placed on grids and stained. 4 μl of the solution were applied onto the grids, most of the protein solution was blotted away with Whatman No.1 filter paper, the grids were washed twice using deionized water, and stained using 1% Uranyl acetate at pH 4.5. After 1 minute, the excess stain was blotted off, and the grids were air dried. Dried grids were stored until imaging at room temperature. Grids were imaged on Philips CM-100 BioTWIN transmission electron microscope with a LaB_6_ electron source at 100 kV (Philips/FEI Corporation, Eindhoven, Holland), fitted with a MegaViewIII digital camera (Olympus Soft Imaging Solutions GmbH, Münster, Germany). iTEM imaging platform (EMSIS GmbH) was used to record micrographs.

### Cryo-EM

MEKK1 TOG–microtubule protein solution was prepared exactly as in the case of NS-EM. 2.5 μl of the solution was applied on glow discharged C-flatTM grids (Protochips) and the grids were blotted and frozen with KF80 specimen plunge-freezing device (Leica) or Vitrobot Mark IV (Thermo Fisher Scientific)

Data collection on frozen-hydrated specimens was carried out using JEOL JEM-2200FS with omega energy filter in cryo configuration with Gatan 914 High-tilt tomography holder and cryotransfer system, fitted with DE-20 4k x 5k direct detector with 6.4 μm x 6.4 μm pixels (Direct Electron). The microscope was operated at 200 kV accelerating voltage with a field emission gun with ZrO/W Schottky emitter. An in-column omega filter with a slit width of 20 eV was used when recording micrographs. The JEOL TEM Controller Application with HT, Gonio and Panel subsystems was used for microscope control and SerialEM was used for semi-automated data collection. 20-frame movies were collected at a dose rate of approximately 1*e*^-^/Å^2^/frame at a magnification of 30000 ×, corresponding to a pixel size of 2 Å. Movie stacks were aligned and dose-weighted using the camera manufacturer’s alignment scripts. Aligned, dose-weighted micrographs were imported to Relion 2 or 3 (He and Scheres, 2017; Nakane et al., 2018). The contrast transfer function (CTF) was estimated using gctf (Zhang, 2016). The CTF fit for each micrograph was examined and poor fits were excluded from further processing. Straight microtubules were manually picked in RELION, avoiding bends, breaks and contamination. Microtubule helix-particles were extracted as overlapping 400×400-pixel particles, offset by 40 px along the central axis of the helix. 2D classification of the extracted particles was performed in RELION, ignoring the CTF until first peak due to the extremely strong structural features of the microtubule lattice distorting the low resolution CTF signature.

### Mammalian cell culture

The HEK293T and COS7 cell lines were cultured in DMEM containing 10% FCS under standard tissue culture conditions (5% CO2, 20% O2). All lines were authenticated by short tandem repeat polymorphism, single-nucleotide polymorphism, and fingerprint analyses, COS-7 cells were seeded in 35 mm imaging dishes coated with 0.8 mg/ml collagen at a density of 2 x 10^5^ cells/dish. 24 hours after seeding, cells were transfected with 1.5 μg of either MEKK1-V5 or MEKK1 V647D/R851E-V5 using Jet Prime transfection reagent. 24 hours after transfection, cell monolayers were scored as in a standard wound healing assay, washed with complete medium and incubated at 37°C for 5 h. Samples were fixed for 20 min with ice cold methanol and stored overnight in PBS at 4°C. Samples were then blocked in 2% BSA/PBS for 30 mins at RT, incubated with rabbit monoclonal anti-V5 tag (CST #13202, 1:500 dilution) and mouse monoclonal anti-β-tubulin (Santa Cruz #sc-58880, 1:200 dilution) primary antibodies for 1 h at RT, washed twice in blocking solution, incubated with goat-anti-rabbit IgG AlexaFluor 488 and goat-anti-mouse IgG AlexaFluor 647 (Jackson ImmunoResearch #111-545-008 and #115-605-166, both 1:250 dilution) secondary antibodies for 30 min at RT, washed twice with PBS, counterstained with DAPI for 5 min at RT and washed twice with PBS. Images were collected with a Leica SP8 scanning confocal microscope using a 63x/1.4 NA objective.

Images of cells displaying a clear polarized morphology were analyzed with ImageJ software. A 90° angle was positioned at the centrosome to define the region of the leading edge that would be analysed. The leading edge was manually demarcated using the MEKK1 signal and a 15-pixel ROI was defined from this line. The coverage of microtubules within this ROI was calculated using an automatic threshold. Once all data were collected, the average threshold was calculated from all WT and MEKK1 images, and all images were reanalyzed using this value. Results were plotted as the percentage of leading edge area containing microtubule signal. The results were checked for correlation between the size of ROI and percentage of leading edge area covered by microtubules.

HEK293T cells were seeded into 6-well plates at a density of 2 x 10^5^ cells/well. 24 hours after seeding cells were transfected with 1 μg of wildtype MEKK1-V5 or 2 μg of mutant MEKK1-V5 using Jet Prime transfection reagent. 24 hours after transfection, lysates were harvested for Western blotting as previously described (Kennedy et al., 2019).

## Supplementary information

**Supplementary Figure 1:**
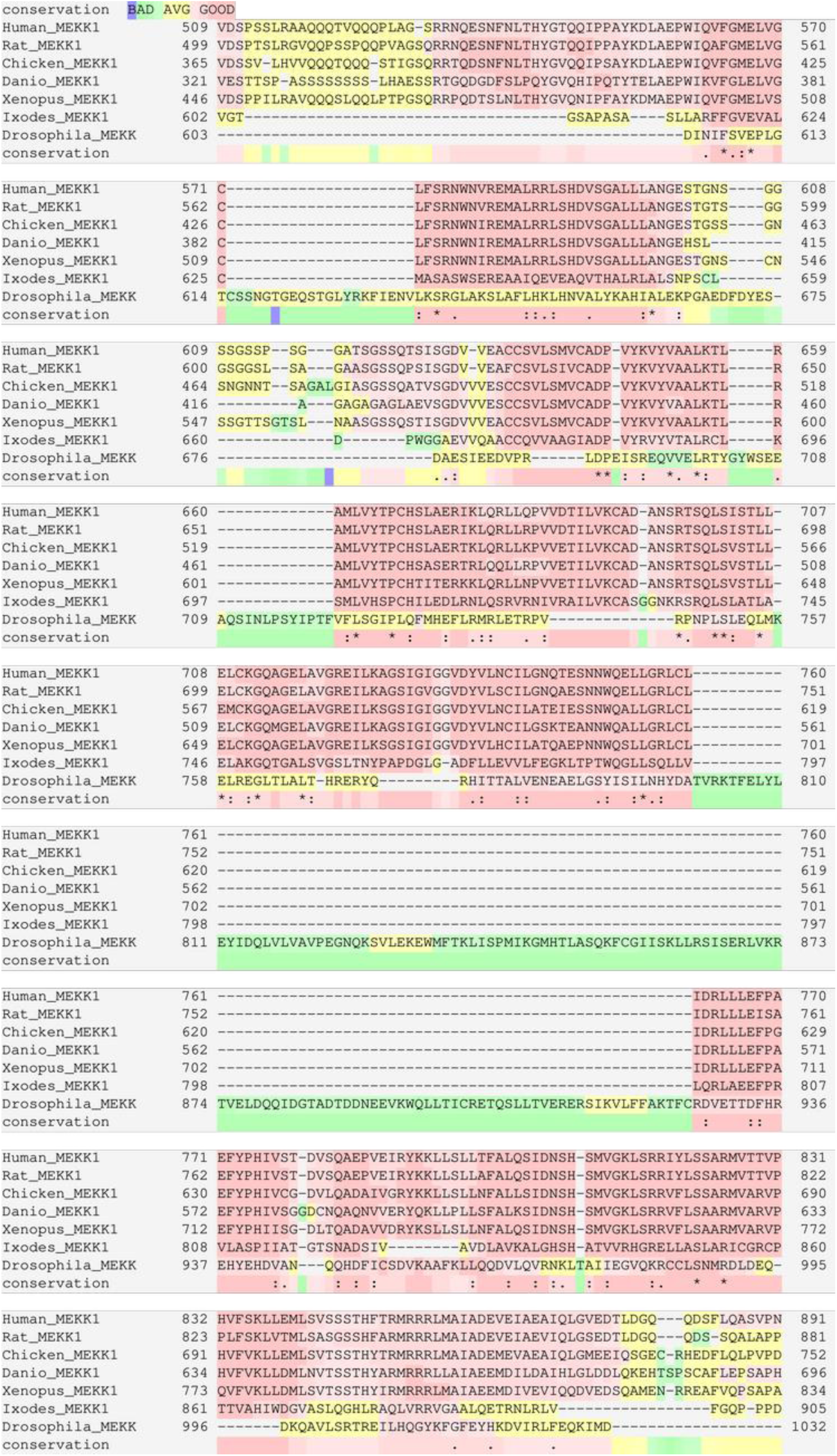
Multiple sequence alignment of MEKK1 (509–891), including the cryptic TOG domain (548– 867) from Human, Rat, Chicken, Zebrafish, Frog, Tick, and Drosophila species. Vertebrates display very high conservation across the TOG domain, Ixodes sp. shows intermediate conservation and Drosophila shows poor alignment, suggesting the domain is functionally lost in Drosophila.

**Supplementary Figure 2:**
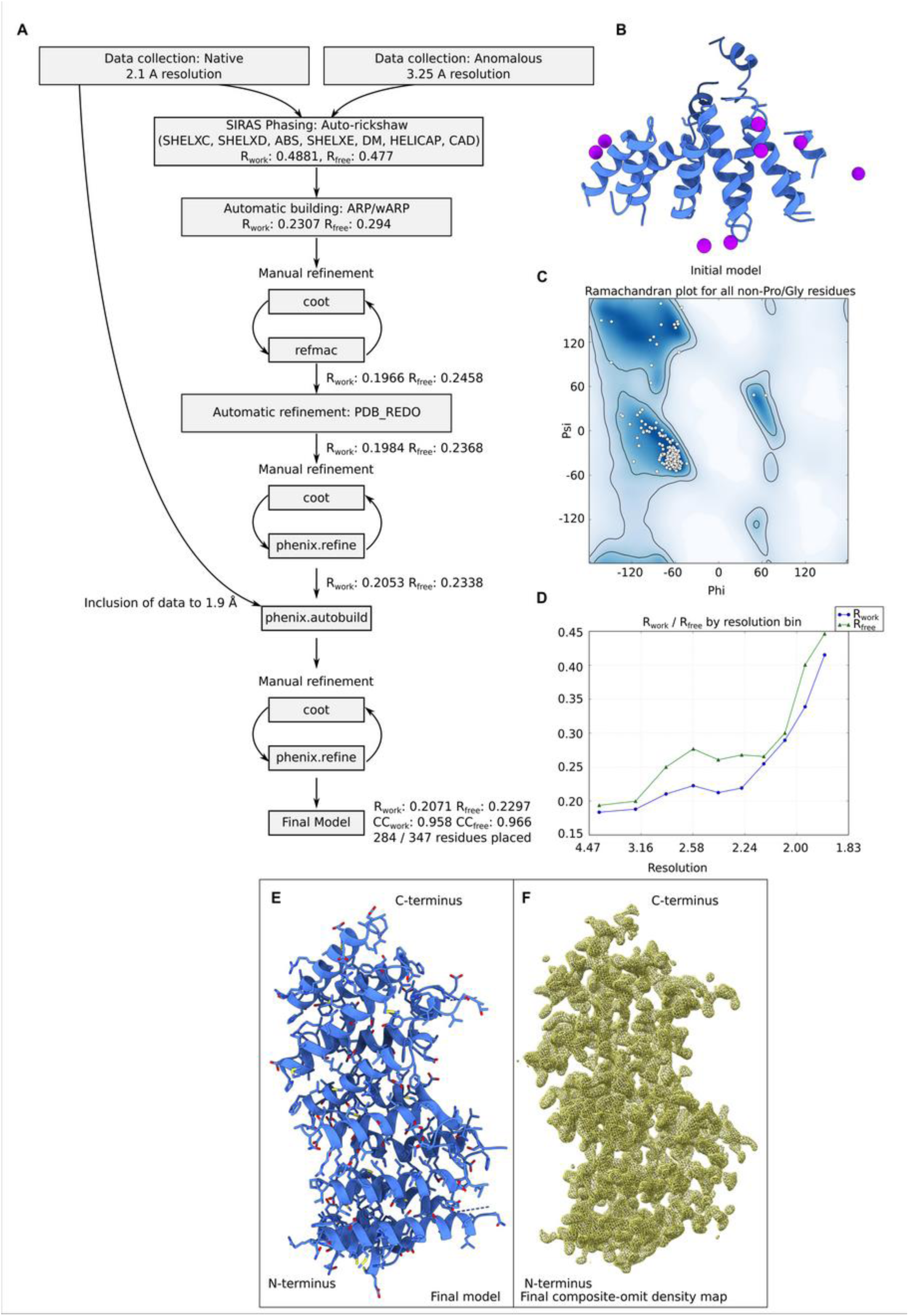
The structure solution and refinement process of MEKK1 central domain. A.) A schematic of structure solution and refinement process. The software used in each step is indicated. B.) Initial phasing model, as output from the Auto-Rickshaw pipeline. Iodine atoms found in the data are shown in purple. The polyalanine model used for subsequent refinement shown in ribbon representation in blue. C.) Ramachandran plot for all non-Pro/Gly residues in the final model shows all residues clustering within the favoured and allowed zones of Psi/Phi space. D.) R-values of the refined model, plotted by resolution bin. E.) The final refined model of MEKK1 TOG domain in blue. F.) A composite-omit electron density map, contoured at 1.5 σ.

**Supplementary Table 1:** Data collection and refinement statistics.

| **MEKK1 TOG domain** | **PDB ID: 6WHB** |
| --- | --- |
| **Wavelength** | 0.9537 Å |
| **Resolution range** | 47.29–1.9 Å (1.968–1.9 Å) |
| **Space group** | I 4<sub>1</sub> 2 2 |
| **Unit cell** | 71.817 Å, 71.817 Å, 259.561 Å, 90°, 90°, 90° |
| **Total reflections** | 782718 (77780) |
| **Unique reflections** | 27403 (2683) |
| **Multiplicity** | 28.6 (29.0) |
| **Completeness** | 99 % (100 %) |
| **Mean I/sigma(I)** | 19.36 (0.55) |
| **Wilson B-factor** | 44.17 |
| **R<sub>merge</sub>** | 0.1447 (8.521) |
| **R<sub>meas</sub>** | 0.1473 (8.669) |
| **CC1/2** | 1 (0.239) |
| **CC\*** | 1 (0.621) |
| **Reflections used in refinement** | 27069 (2356) |
| **Reflections used for R-free** | 1381 (113) |
| **R<sub>work</sub>** | 0.2071 (0.4153) |
| **R<sub>free</sub>** | 0.2297 (0.4466) |
| **CC<sub>(work)</sub>** | 0.958 (0.492) |
| **CC<sub>(free)</sub>** | 0.966 (0.497) |
| **Number of non-hydrogen atoms** | 2326 |
| **macromolecules** | 2220 |
| **ligands** | 38 |
| **Protein residues** | 284 |
| **RMS(bonds)** | 0.006 Å |
| **RMS(angles)** | 0.73° |
| **Ramachandran favored** | 99 % |
| **Ramachandran allowed** | 1.1 % |
| **Ramachandran outliers** | 0 % |
| **Rotamer outliers** | 0.4 % |
| **Clashscore** | 1.31 |
| **Average B-factor** | 55.59 Å<sup>2</sup> |
| **macromolecules** | 55.09 Å<sup>2</sup> |
| **ligands** | 87.08 Å<sup>2</sup> |
| **solvent** | 54.50 Å<sup>2</sup> |
| **Number of TLS groups** | 1 |

Statistics for the highest-resolution shell are shown in parentheses.

**Supplementary Figure 3:**
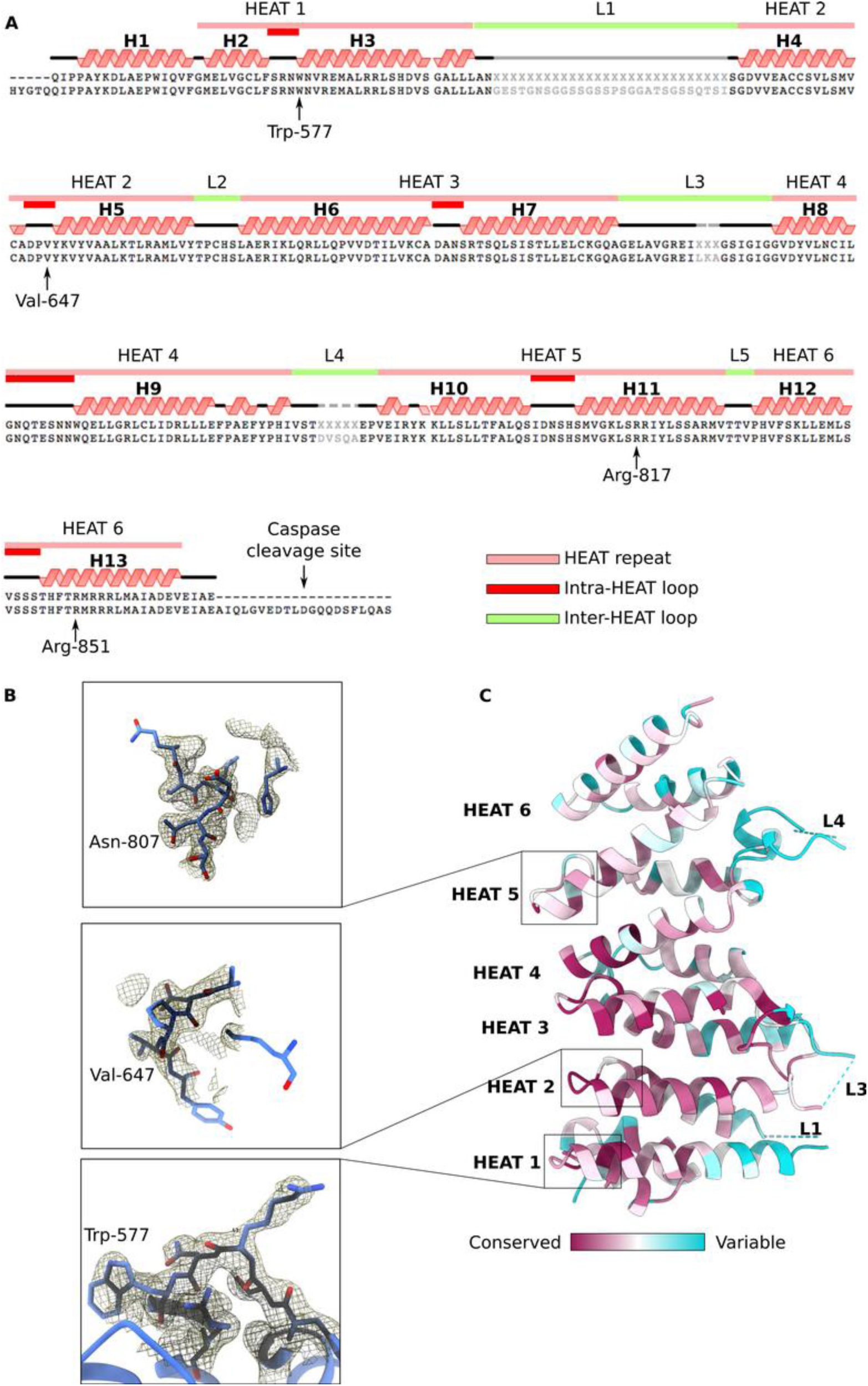
Sequence, structure and conservation analysis of MEKK1 TOG domain. A.) Secondary structure elements defined in the crystal structure of MEKK1 TOG. Helices are numbered for the N-terminus as H1-H13. Schematic view of HEAT repeats, intra- and inter-HEAT loops on top, secondary structure assignment in the middle, and alignment of the structured sequence with the crystallization construct sequence on the bottom. B.) Electron density definition of intra-HEAT 1, 2, and 5 loops in composite-omit map, contoured at 1.5 σ. C.) ConSurf analysis of the MEKK1 TOG domain reveals high conservation in the N-terminal HEAT repeats and in the intra-HEAT loops on the tubulin-interacting side, and low conservation on the opposite side containing long, disordered loops.

**Supplementary Table 2:** PDBeFold search for structural homologs of the crystal structure of the central domain of MEKK1 revealed structural similarity to TOG domains.

| ## | Q-score | RMSD | Alignment |  |  |  | PDB ID |
| --- | --- | --- | --- | --- | --- | --- | --- |
|  |  |  | Nalgn | Nsse | Ngaps | Seq-% |  |
| 1 | 0.37 | 2.682 | 220 | 12 | 8 | 0.1955 | 5jo8:A Cep104 TOG |
| 2 | 0.3683 | 2.665 | 221 | 12 | 11 | 0.1538 | 5lph:B Cep104 TOG |
| 3 | 0.3668 | 2.743 | 223 | 12 | 10 | 0.1749 | 5lph:A Cep104 TOG |
| 4 | 0.3003 | 3.066 | 202 | 11 | 12 | 0.1436 | 6mq5:A CLASP1 TOG1 |
| 5 | 0.2836 | 3.012 | 197 | 11 | 9 | 0.1929 | 5dn7:A Crescerin TOG |
| 6 | 0.2353 | 3.34 | 198 | 11 | 12 | 0.1111 | 5vjc:A Msps TOG |
| **MEKK1 TOG** |  |  | **284** | **15** |  |  |  |

**Supplementary Table 3:**
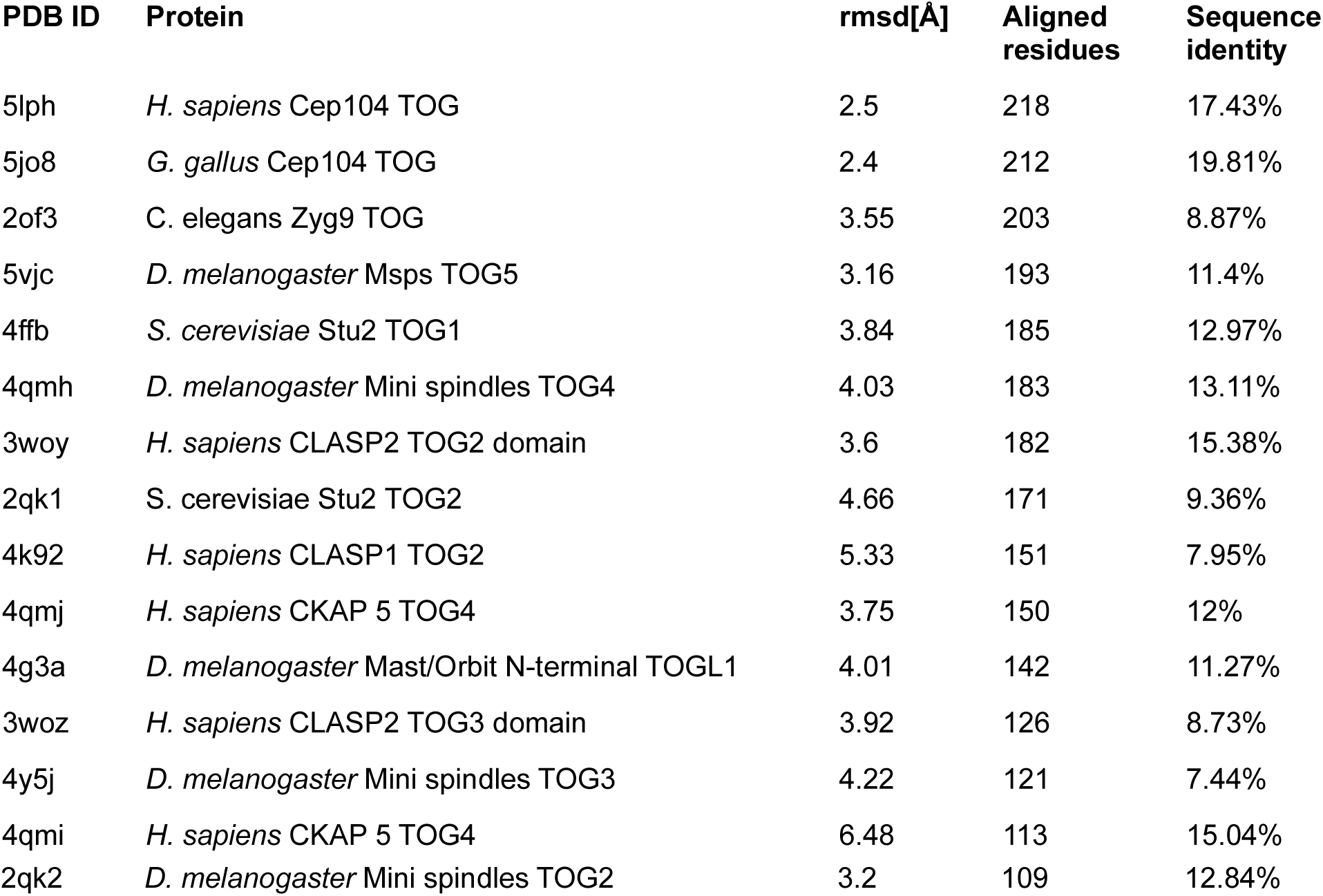
Root-mean-square distances between corresponding Cα atoms of MEKK1 TOG and known structures of other TOG domains, sorted by length of alignment. Secondary structure matching superpositions calculated by coot.

**Supplementary Figure 4:**
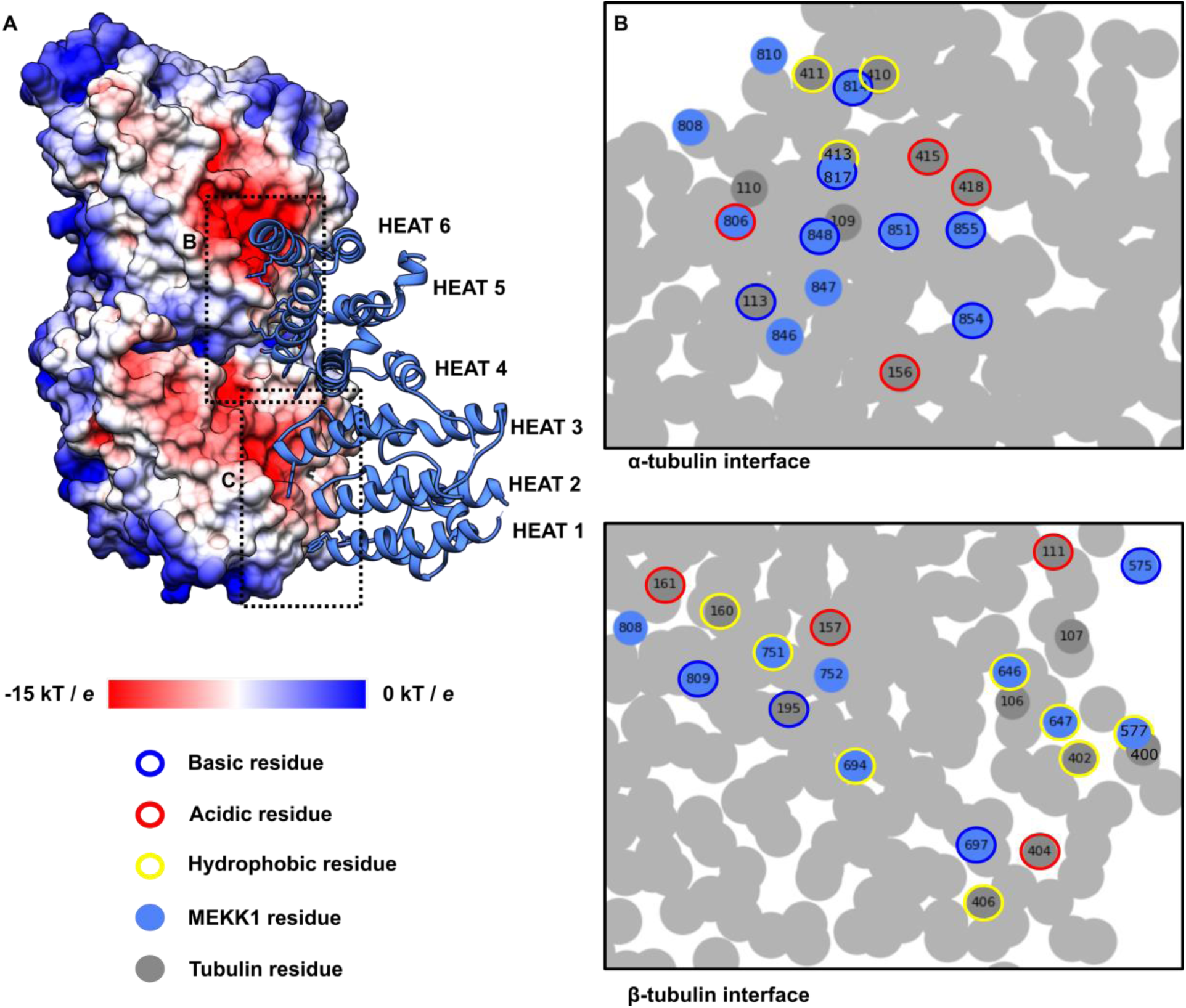
Interface analysis in ChimeraX reveals potential for many interactions between MEKK1 TOG and tubulin. A.) An interaction model of MEKK1 TOG and tubulin prepared by overlaying MEKK1 TOG on Stu2p TOG1 in complex with tubulin (PBD ID: 4FFB). B.) An interaction footprint of MEKK1 TOG on α-tubulin. C.) An interaction footprint of MEKK1 TOG on β-tubulin.

**Supplementary Figure 5:**
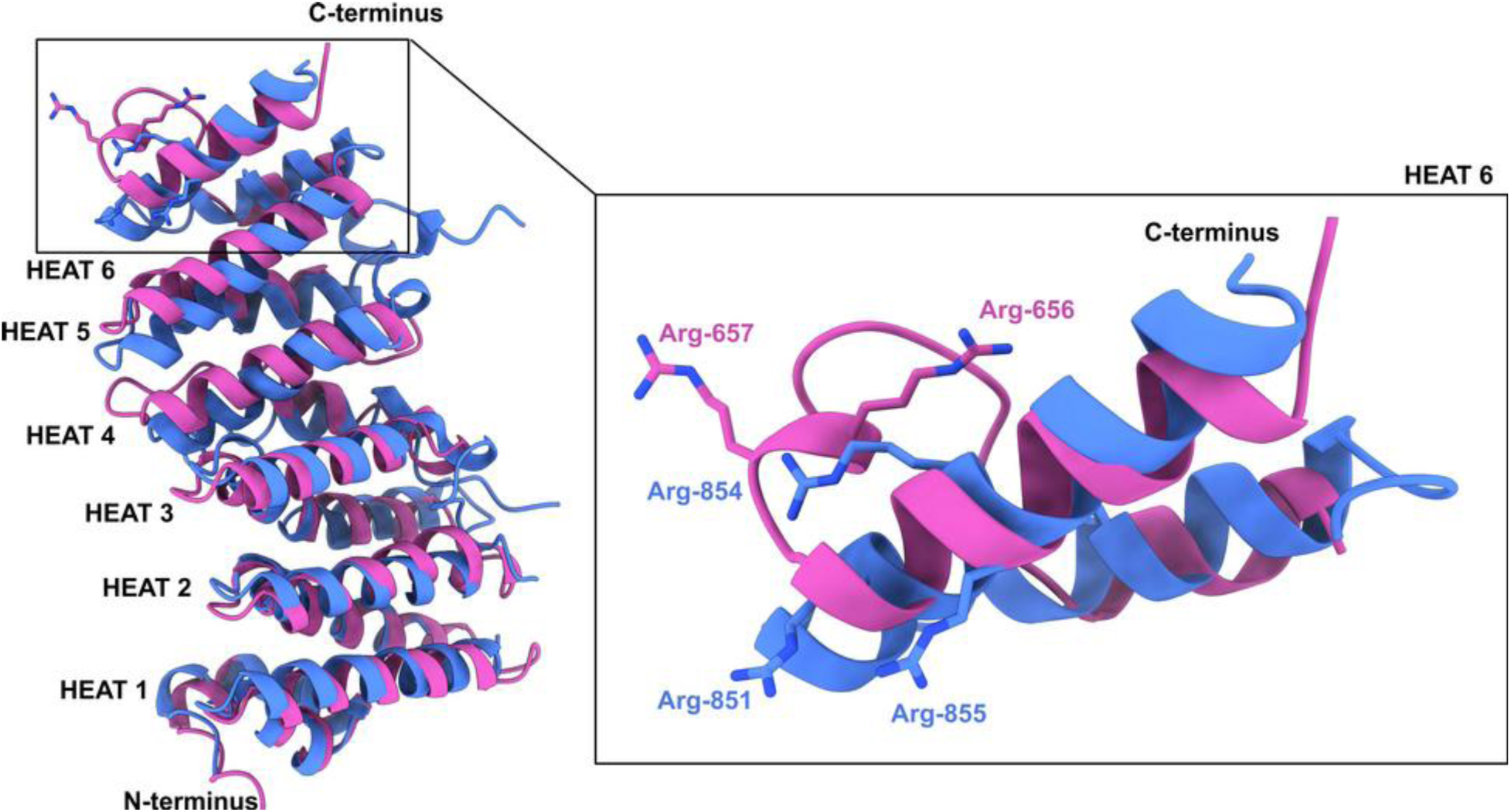
A cartoon overlay of MEKK1 TOG (blue) and Cep104 TOG (pink, pdb:5jo8) domains with C-terminal arginines shown as sticks. A.) A full overlay, showing good structural conservation across the whole structure, including the N-terminal extension of MEKK1 TOG. The C-termini are both arginine-rich, but differ structurally. B.) A detailed view of the C-terminal HEAT repeat of MEKK1 and Cep104 TOG domains.

**Supplementary Figure 6:**
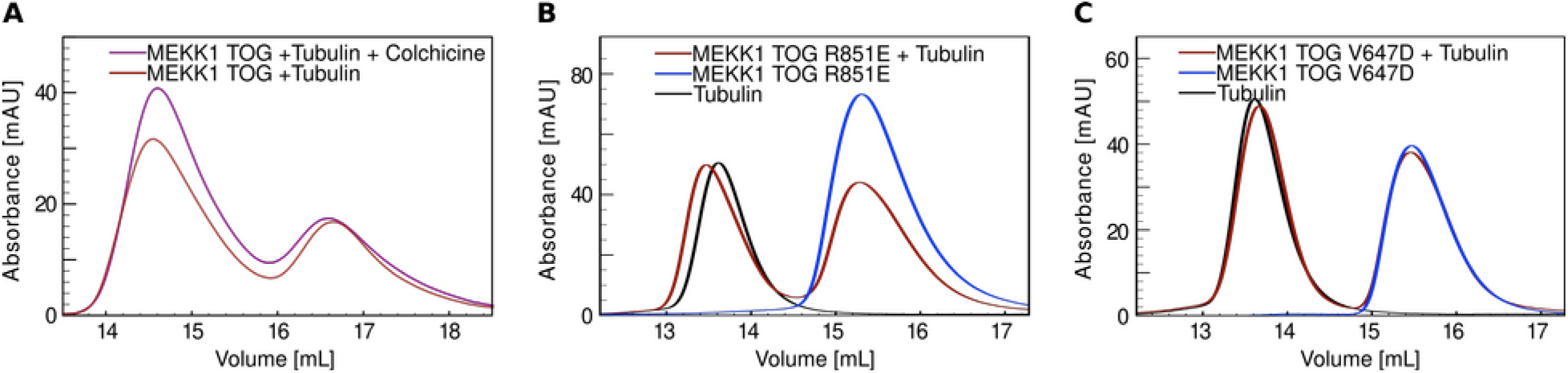
Solution analysis of colchicine-treated tubulin binding by MEKK1 TOG wt and free tubulin binding by MEKK1 mutants. A.) SEC trace of wt MEKK1 TOG binding to free (red) and colchicine-treated (purple) tubulin. The elution volume shift is negligible. B.) SEC trace of MEKK1 TOG R851E (blue), free tubulin (black), and their combination (red). C.) SEC trace of MEKK1 TOG V647D (blue), free tubulin (black), and their combination (red).

**Supplementary Figure 7:**
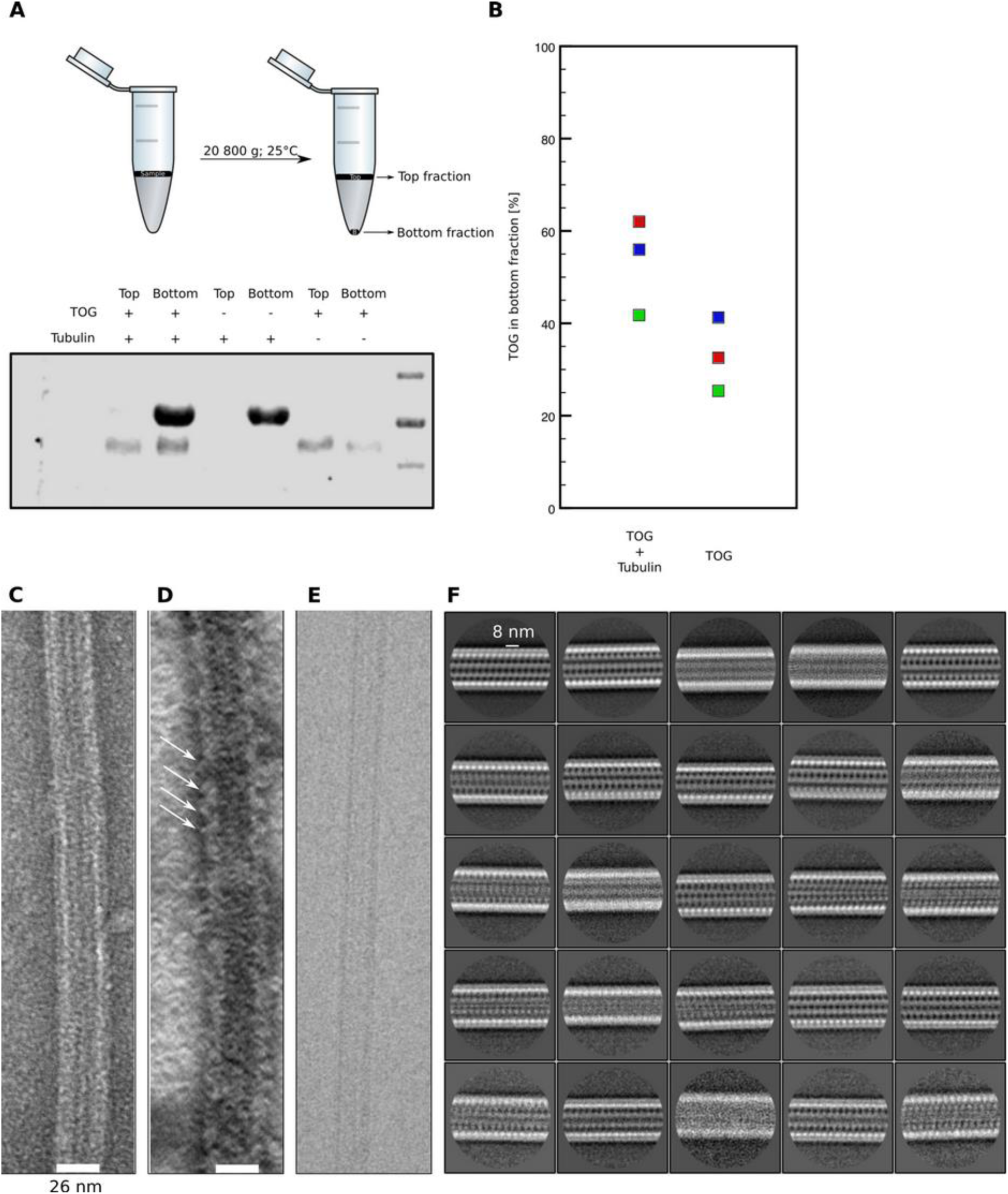
The assessment of the capacity of MEKK1 TOG to bind to the straight conformation of tubulin. A.) Cosedimentation assays hinted as the possibility of MEKK1 TOG binding to microtubules. MEKK1 TOG appeared in the bottom fraction in greater quantity in presence of tubulin than in absence of tubulin. B.) A quantification of three replicate cosedimentation assays. C.) NS-EM of a paclitaxel-stabilized microtubule. D.) NS-EM of a paclitaxel stabilized microtubule with MEKK1 TOG, showing protein densities binding to the outside of the microtubule in regular intervals. E.) An aligned, dose-weighted cryo-EM micrograph of a microtubule in presence of MEKK1 TOG, collected with 2 μm defocus. F.) A 2D classification of microtubules in presence of MEKK1 TOG. The class averages show no densities consistent with the binding of MEKK1 TOG on the outside of the microtubule.

**Supplementary Figure 8:**
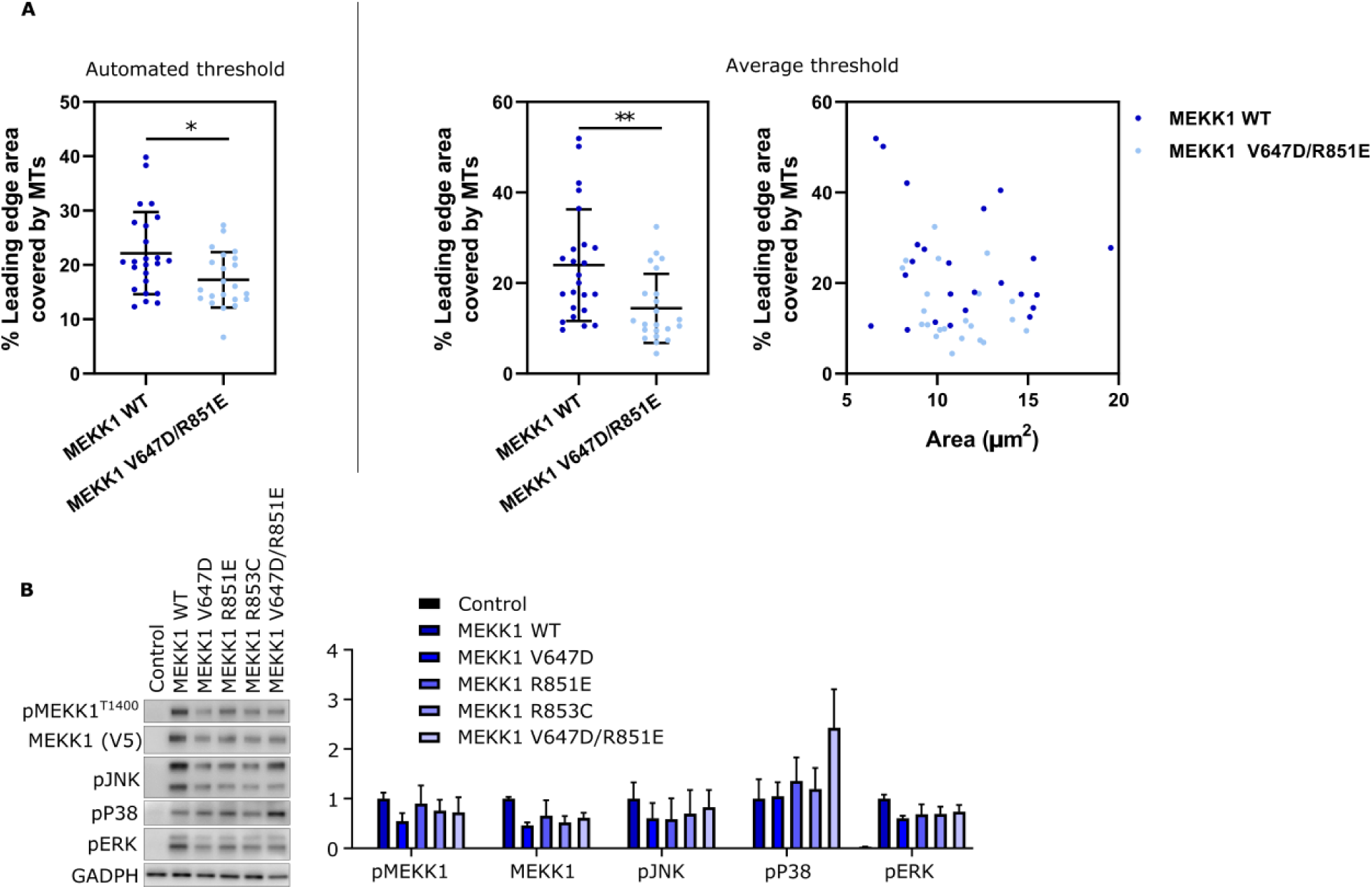
The analysis of the leading edge area and the analysis of MAPK activation. The analysis of the leading edge area was performed using both individual (A., left panel) and average (A., middle panel) thresholding approach. The proportion of leading edge area covered by microtubules was independent of the total area demarcated as leading edge (A., right panel). B.) Analysis of mutant MEKK1 in HEK-293T cells. Left panel: A representative western blot analysis of the activation of downstream MAPK pathways upon overexpression of wt and mutant MEKK1. Right panel: A quantification of western blot analyses, as shown on the left, from three biological replicates.

